# Combining computational biology and experimental knowledge to draw a structural profile of the active membrane-assembled NADPH oxidase complex

**DOI:** 10.1101/2024.02.19.579638

**Authors:** Sana Aimeur, Burcu Aykac Fas, Xavier Serfaty, Hubert Santuz, Sophie Sacquin-Mora, Tania Bizouarn, Antoine Taly, Laura Baciou

## Abstract

The phagocyte NADPH oxidase complex (NOX2) is activated by various intracellular stimuli, resulting in the generation of reactive oxygen species, which are responsible for many physiological processes, including innate immune defense. Structures of the cytochrome *b*_558_, membrane catalytic core of the complex, in their resting state have been solved experimentally. Unfortunately, very little is known about the structure of the active complex, as it involves the assembly of modular proteins and disordered regions, that have been shown to be essential to the function of the enzyme. Here, we modelled the supra-structure of the active complex using Alphafold2 and taking into account experimental data as well as conformational analyses by SRCD. AF2 ability to generate complexes enabled us to visualize ways in which subunits could assemble together, and revealed the implications of intrinsically disordered regions in protein-protein and membrane-protein interactions. Additional Molecular Dynamics simulations of the membrane embedded complex bring further insight regarding the system dynamics. Altogether, our work provides evidence for a model of the structure of the active complex where disordered regions can enable multi-subunit assembly.

## Introduction

Proteins during their function can undergo many conformational changes related to post-translational modifications but also related to interactions with other molecules, proteins or lipid membranes with which they can interact. This is the case of the NADPH oxidase which is dedicated to the production of superoxide anion (O_2 -_), precursor of reactive oxygen species (1). In phagocyte cells, NADPH oxidase-generated ROS cause the destruction of invading pathogens. The minimal functional phagocyte NADPH oxidase complex is composed of five subunits: the flavocytochrome *b*_558_ (cyt*b*_558_), a membrane-bound heterodimer which consists of NOX2 (initially named gp91^phox^) and p22^phox^, and three cytosolic subunits p47^phox^, p67^phox^ and a G-protein of the Rho family (Rac1 or Rac2). The cyt*b*_558_ activity is dependent on the assembly with the cytosolic proteins on its cytosolic side. When the enzyme is down-regulated, it leads to severe immune deficiency as illustrated by the chronic granulomatous disease (CGD) while when it is up-regulated, it participates to inflammatory and oxidative stress-related diseases. Thus, fine-tuning the multi-component assembly is essential to regulate enzyme functioning by avoiding aberrant levels of ROS production and generating specific reactive oxygen species when and where they are needed.

Once the complex is assembled, the catalytic subunit NOX2, that carries all redox carriers, is able to transfer electrons across the membrane from the substrate, NADPH, to molecular oxygen to produce O_2 -_. Two hemes are located in the N-terminus transmembrane (TM) domain of NOX2. One flavin and the substrate NADPH binding sites are located in the C-terminus dehydrogenase (DH) domain of NOX2, domain which resides at the cytoplasmic face of the membrane. The flavin acts as the initial acceptor of a pair of electrons from NADPH and subsequently transfers one by one the electrons to two molecular oxygen through the hemes. When only associated to p22^phox^, NOX2 is considered in the resting state, unable to bind NADPH and to perform any transmembrane electron transfers. In resting cells, the cytosolic proteins p47^phox^, p67^phox^ and Rac1 or Rac2 are dispersed into the cytosol. In activated cells, in response to soluble phagocyte stimuli or upon pathogen phagocytosis, the docking of the cytosolic subunits on the cytosolic side of the cyt*b*_558_, composed mainly of the DH domain of NOX2 and the intracellular C-terminus of p22^phox^ is triggered by post-translational modifications (PTM) achieved by intracellular signaling cascades. These PTM, including notably the phosphorylations of the oxidase subunits and lipid modifications, are known to initiate, through conformational changes, the protein-protein and lipid-protein interactions essential for the edifice of the functional oligomerized complex (reviewed in (2)). This complex choreography between the different proteins (cytosolic and membrane proteins) must be precisely achieved in phagocytes to result in an active NADPH oxidase complex. The p47^phox^ subunit plays an important role in this mechanism since it mediates the first steps in assembling the active oxidase complex (3). Briefly, phosphorylations induce substantial conformational changes of p47^phox^ which favour its interaction with p22^phox^ and with the membrane. Also known as the “organizer”, it drives the translocation of p67^phox^ to the membrane and to the NOX2 subunit (4).

The recent determination of the cryo-EM structures of cyt*b*_558_ has provided an important molecular view of the membrane subunits in their resting state (PDB code 7U8G and 8GZ3) (5, 6). However, unravelling the structure of the functional (assembled) complex remains a challenge, considering the fact that the assembly of the complex is likely to be a dynamic process occurring step by step (in cascade). Fortunately, several studies have enabled to describe certain aspects of the interactions involved in the complex activation. For example, functional studies have described the crucial interaction between SH3 domains of p47^phox^ to proline-rich sequences in p22^phox^ (7). This was confirmed by structural studies describing the interacting region of p47^phox^ with a peptide of 18 amino acid length from p22^phox^ (8, 9). This C-terminus p22^phox^ peptide from Lys149 to Glu169 contains a Proline Rich Region (PRR) comprising the consensus motif PxxP. This region of the protein was crystallized with the SH3 domain of p47^phox^. This segment is important for the assembly and activation of NOX2. Interactions between specific domains of p67^phox^ and p47^phox^ (10) and between Rac and p67^phox^ (11) have also been described. Nevertheless, this structural information is based only on fragments (peptide) or domains of these proteins. In addition, many other studies were performed to unravel the complex assembly, some of them implying the crucial role of the surrounding membrane, including specific lipids (12–14) (15, 16) or the PX domain of p47^phox^ and the PB region of Rac described as being essential to the complex assembly. All this information was used to propose a scenario for assembling the complex but the overall structure of the active complex, and the structural changes required of the various protein partners to form it remain elusive. This is hampering the search for molecule regulators for potential therapeutic development targeting the functional enzyme.

In this work, we took advantage of the rapid development of computational biology concerning the *in silico* prediction of protein structures. We combined the structures available in the AlphaFold2 (AF2) database with the prediction of multi-subunit complexes using AF2 based on protein-by-protein interaction to map the active complex architecture. This enabled us to explore the network architecture of the assembled NADPH oxidase complex inserted into predicted membrane. In addition, we ran all-atom classical Molecular Dynamics simulations on the membrane embedded complex, which brought us further information regarding the system dynamic properties, and in particular, the stability of the contacts formed between the different domains. Finally, in order to make the best possible progress towards the optimal profile of the active state of the NADPH oxidase complex, we have drawn on spectroscopic data and knowledge from the literature.

## Materials and methods

### Materials

Chemicals (*ci*s-arachidonic acid, phosphate buffered saline (PBS)) came from Sigma Aldrich (ST Quentin Fallavier, France). Resins Ni-sepharose, SP-sepharose HP were purchased from Cytiva. Cis Arachidonic acid (AA) was solubilized at a concentration of 20 mg/mL in ethanol, aliquoted and stored at −80°C.

### In vitro studies

#### Expression and purification of Trimera

The Trimera plasmid was generously donated by Prof. E. Pick (Tel Aviv University, Israel). Trimera is a chimeric protein consisting of the fusion of human N-part of p47^phox^ (aa 1-286), N-part of p67^phox^ (1–212) and full-length Rac1Q61L (1–192) (17). Expression and purification of trimers were carried out as previously described in (17). Purification chromatography including Ni-NTA-Sepharose-FF resins (Cityva, France) was carried out using ÄKTAprime system. The purity of protein solutions was assessed by migration in 10 % SDS BisTris-NuPAGE gels (Biorad), stained with Coomassie Brilliant Blue. Protein concentrations were determined with a NanoDrop^TM^ spectrophotometer at 280 nm using the absorption coefficient calculated for the Trimera (ExPASy, ProtParam free software).

#### Circular dichroism spectroscopy

Circular dichroism (CD) was used to explore the secondary structure of the proteins in solution and the structural changes associated to arachidonic acid-protein complex. The protein samples were dialyzed overnight in 100 mM NaF and 10 mM of NaPO_4_ (pH 7.0). The dialysate buffer was retained for CD control analyses. Prior to data collection, samples were briefly centrifuged at 10,000 x g to remove protein aggregates.

Synchrotron radiation circular dichroism (SRCD) spectra were collected on the DISCO beamline at the SOLEIL synchrotron radiation facility, Gif-sur-Yvette, France(18, 19). The use of synchrotron radiation enables us to extend the spectral range to lower wavelengths than would be possible with classical laboratory equipment. All spectra were calibrated for wavelengths and amplitudes with camphor sulphonic acid (CSA). Protein samples ( 3.6 mg/mL) were loaded in CaF_2_ cells using 3 µL volumes and optical pathlengths of 50 µm (19). When indicated, arachidonic acid was added as an ethanolic solution as it is not soluble in water. An identical volume of ethanol was added in the reference, in order to subtract its CD contribution. We have checked that ethanol at such concentration did not perturb the secondary structure of the proteins.

Spectra were measured from 280 to 170 nm, using the HT (high tension) mid-eight as the cut-off at 175 nm. Three consecutive scans were recorded and averaged for the samples and the baseline. The mean baseline was subtracted from the mean sample spectra. The temperature was kept constant at 25°C.

Spectra are expressed in delta epsilon units, calculated using mean residue weight of Trimera (pending for 730 amino acid length). The secondary structure composition was determined using the BestSel software. The fitting range was 180-250 nm.

## Modelling approaches

### In silico protein complex modelling

The up-graded version of the AlphaFold 2 (AF 2.3) algorithm was used to predict the three-dimensional structures of NADPH oxidase protein complexes from their amino acid sequences (https://alphafold.ebi.ac.uk/) (20, 21). It was used online or implemented locally to increase the number of cycles. The number of cycles was set at 18, as no further improvements were observed in the models generated beyond this number.

In silico predicted protein circular dichroism spectra were performed using the SESCA software(22). A method for Positioning of Proteins in Membrane (PPM2.0) was used to model the cyt*b*_558_ orientation in a lipid bilayer using the PPM server (https://opm.phar.umich.edu/ppm_server) (23). The membrane interface or “midpolar region” (8-20 Å from the membrane center) is characterized by the steep gradient of all polarity parameters whose values depend on lipid composition fixed as plasma membrane from mammals (**Table 1**).

**Table 1:**
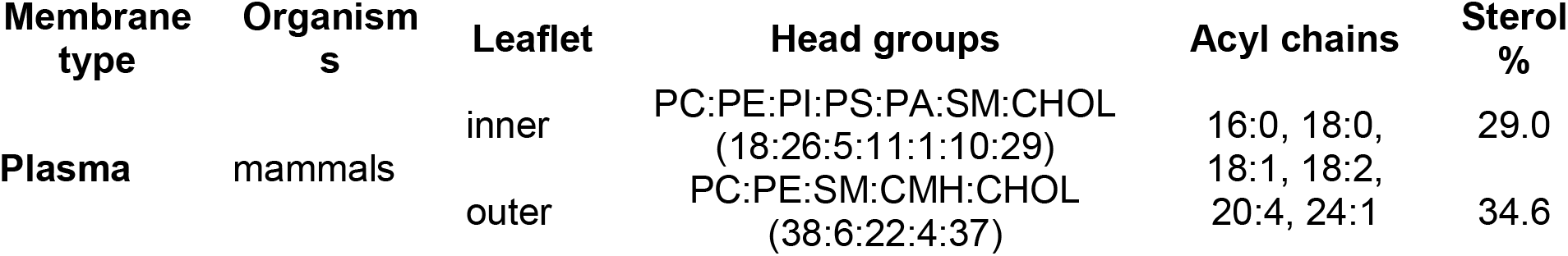
Complex lipid compositions of simulated plasma membrane using PPM web server (https://opm.phar.umich.edu)

### *In silico* protein domain movement analysis

The DynDom server (http://dyndom.cmp.uea.ac.uk/dyndom/) was used to determine the angle of deviation of protein domains from the membrane by displaying domain movements relative to an axis of rotation (24). The coordinates of the different structural models were used as input data to obtain the angle of deviation of the DH domain from the membrane.

### *In silico* cofactor insertion

The AlphaFill algorithm (https://alphafill.eu/) was used to insert the cofactors into the predicted structures(25). The prediction is based on at least 25 % sequence identity with the reference protein, and RMSD metrics are calculated on the basis of residues within 6 Å around the cofactor.

### Molecular Dynamics

In this study, molecular dynamics simulations of the NOX2-p22^phox^-p47^phox^ complex embedded within a membrane bilayer were conducted using GROMACS version 2021.3 to explore the dynamics (26, 27). The initial structure, which was modeled as described in the *In silico protein complex modeling* section, was further processed with the Phenix software (28) to minimize clashes and strains. The model was then oriented for membrane embedding using the PPM server (29).

The NOX2 protein was modified with glycans at residues N132, N149, and N240, and featured a disulfide bond between C244 and C257, in addition to heme coordination centers at outer heme/heme2 (H115-H222) and inner heme/Heme1 (H101-H209). The membrane system, assembled using CHARMM-GUI (24, 30, 31), mimicked the plasma membrane with a lipid composition of POPC, PLPC, DOPC, POPE, PLPE, DOPE, POPS, cholesterol, PI(3)P, and PI(3,4)P2 (**Table 2**). Simulations were performed in an explicit TIP3P water model (32), within a hexagonal periodic box (**Table 3**). The systems underwent energy minimization, NVT, and NPT ensemble equilibrations, ensuring stabilization before the production phase. We conducted two 500 ns simulations for the NOX2-p22^phox^-p47^phox^ complex. The phosphorylated form of the p47^phox^ protein was used with phosphorylations introduced at S303, S304, S328, S359, S370, and S379 residues. The simulations were performed using the CHARMM36m force field, recognized for its accuracy in modeling protein-lipid interactions (33), with a time step of 0.002 ps. Long-range electrostatic interactions were treated using the Particle Mesh Ewald (PME) method (34), with van der Waals interactions managed using a cut-off and force-switch modifier at 1.0 and 1.2 nm, respectively.

**Table 2:** Lipid composition used in the all-atom molecular dynamics simulations to mimic the plasma membrane.

| Lipid Type | Lipid Name | Outer Leaflet | Inner Leaflet |
| --- | --- | --- | --- |
| CHL1 | Cholesterol | 93 | 52 |
| POPC | 1-Palmitoyl-2-oleoylphosphatidylcholine<br>(C<sub>42</sub>H<sub>82</sub>NO<sub>8</sub>P) | 30 | 26 |
| PLPC | 1-Palmitoyl-2-linoleoyl-sn-glycero-3-phosphatidylcholine<br>(C<sub>42</sub>H<sub>80</sub>NO<sub>8</sub>P) | 50 | 33 |
| DOPC | Dioleoyl phosphatidylcholine<br>(C<sub>44</sub>H<sub>84</sub>NO<sub>8</sub>P) | 15 | 18 |
| POPE | 1-Palmitoyl-2-oleoylphosphatidylethanolamine<br>(C<sub>39</sub>H<sub>76</sub>NO<sub>8</sub>P) | 30 | 29 |
| PLPE | 1-Palmitoyl-2-linoleoyl-sn-glycero-3-phosphatidylethanolamine<br>(C<sub>39</sub>H<sub>74</sub>NO<sub>8</sub>P) | 30 | 28 |
| **DOPE** | **1,2-Dioleoyl-sn-glycero-3-phosphoethanolamine (C<sub>41</sub>H<sub>78</sub>NO<sub>8</sub>P)** | **16** | **30** |
| **POPS** | **1-Palmitoyl-2-oleoylglycero-3-phosphoserine (C<sub>40</sub>H<sub>76</sub>NO<sub>10</sub>P)** | **0** | **12** |
| **SAPI13** | **Phosphatidylinositol 3-phosphate (PI(3)P)** | **0** | **2** |
| **SAPI2C, SAPI2D** | **Phosphatidylinositol 3,4-bisphosphate (PI(3,4)P<sub>2</sub>)** | **0** | **2** |
| **TOTAL LIPIDS** |  | **264** | **232** |

**Table 3:** Details of the simulated systems.

| **Simulated System** | **No. of residues** | **Box dimension AxBxC (Å)** | **Total no. of atoms** | **No. of water molecules** | **No. of Lipids: Upper/Lower Leaflet** | **No. of K+/Cl- ions (Concentration)** | **Simulation time (ns)** | **Replicas** |
| --- | --- | --- | --- | --- | --- | --- | --- | --- |
| NOX2-p22phox-p47phox | 1155 | 136x136x212 | 325032 | 83146 | 264/232 | 245/228 (0.15 M) | 500 | 2 |

*\* All simulations of the NOX2-p22phox-p47phox model were conducted within an hexagonal box under periodic boundary conditions at 310.15 K. The p47phox subunit was phosphorylated at residues S303, S304, S328, S359, S370, and S379. Interactions were modeled using the CHARMM36m force field with the TIP3P water model, conducted on Gromacs 2021.3.*

Temperature and pressure were regulated using the Nose-Hoover thermostat and the Parrinello-Rahman barostat, set at 310.15 K and 1.0 bar. Hydrogen bond constraints were applied using the LINCS algorithm. The model included hemes and FAD cofactors, with FAD parameters sourced from (35). The placement of the hemes was taken from the cryo-EM structure 7U8G (6) and FAD was modeled based on the DUOX1-DUOXA1 cryo-EM structure 7D3E (36). The DH domain of the model was shifted to match the orientation of the DH domain of 7D3E and to adjust the distance between the inner heme and FAD, to serve as a physically relevant starting point for MD simulations, which is expected to further refine the model.

## Results

### AF2-generated cytb_558_ model and comparison with the experimental resting cytb_558_ structure

AF2 was used to generate the structural model of cyt*b*_558_ consisting in the heterodimer of NOX2 and p22^phox^. For this purpose, the human NOX2 and p22^phox^ sequences were executed in the AF2 input parameters. The five cyt*b*_558_ models generated by AF2 showed overlapped membrane regions of NOX2 and p22^phox^, differences lying mainly in the more flexible and disordered cytosolic regions of NOX2 and p22^phox^ (**Figure 1A**).

**Figure 1.**
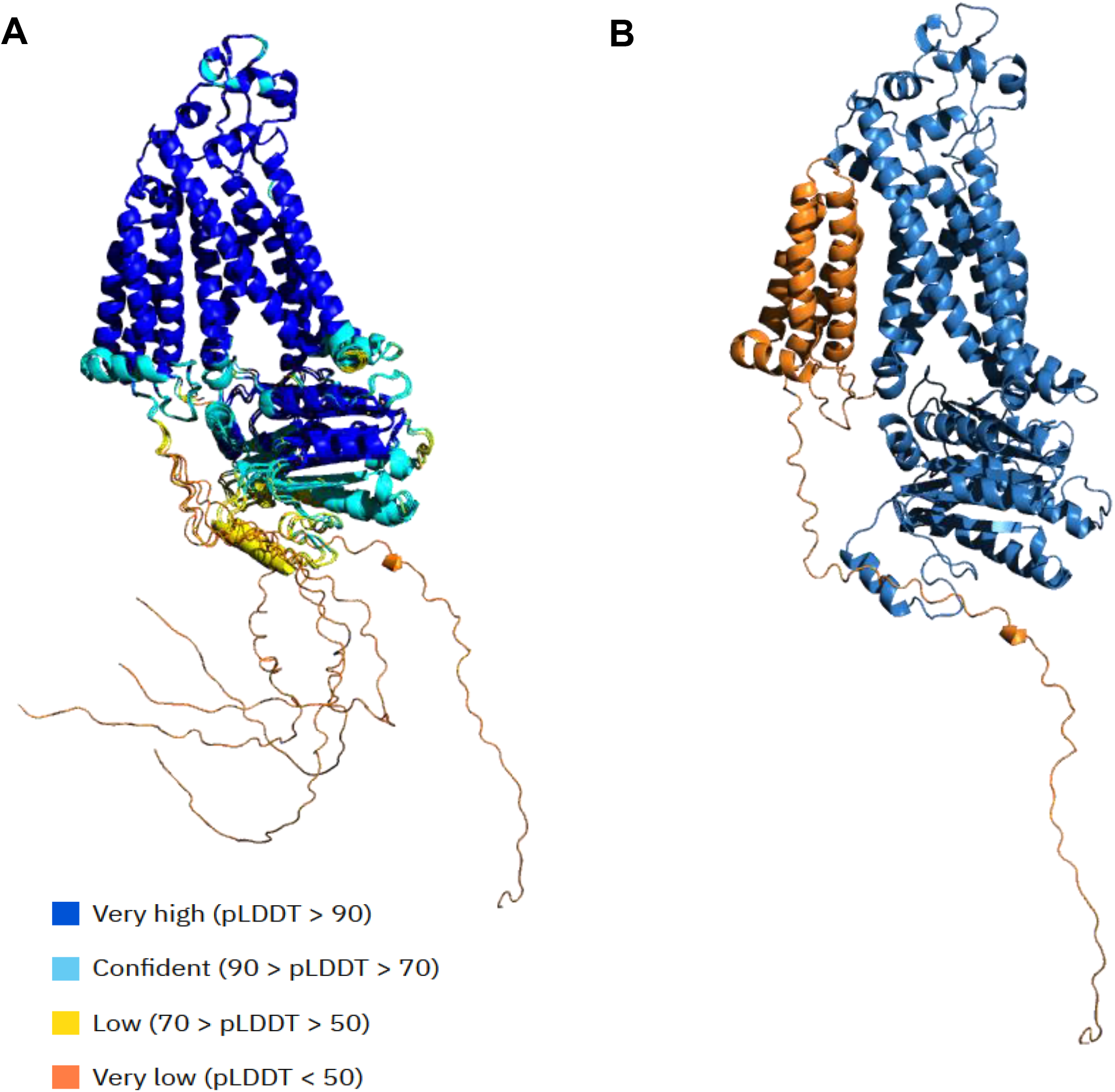
The AlphaFold2-generated Cyt*b*_558_ models. **A. Superposition of the five models of cyt*b*_558_ generated by AF2 locally implemented.** Alignment was performed with PyMOL software based on the membrane region of NOX2. The colors describe the pLLDT score, showing the confidence levels of the different parts of the structure. B. Selected AF2 Cyt*b*_558_ model. NOX2 is colored in blue and p22^phox^ is shown in orange.

Consistently with the recent experimental structures of cyt*b*_558_ in resting state (7U8G and 8GZ3) that was published in 2022 after the latest retraining of AF2 (the data cutoff of version 2.3 was in 2021), the predicted NOX2 is organized in 6 transmembrane helices (TM) (**Figure 1B**). Its superposition with 7U8G and 8GZ3 gave a RMSD of 0.446 Å and 0.405 Å, respectively (5, 6). The predicted p22^phox^ is organized in 4 transmembrane helices with the C-and N-terminal regions oriented towards the intracellular side (**Figure 1B**), as proposed first by the team of E. Pick (37) based on peptide walking studies, prior to any structural data. It should be noted that the p22^phox^ topology was subject of several papers disagreeing on the number of TM helices. Furthermore, the 4-helix transmembrane organization of p22^phox^ was proposed by AF2 prior to any structural experimental release. In the AF2 models of cyt*b*_558_, the C-terminal region of p22^phox^ remains entirely disordered and shows no identifiable structure except a small -helix located at the end of the PRR region of p22^phox^ (**Figure 1A and B**). The interaction between NOX2 and p22^phox^ in the AF2 model is similar to that described in the experimental structural model (**Figure 2A**) and imply the same residues (**Figure S1**) demonstrating the accuracy of membrane protein complex prediction by AF2.

**Figure 2.**
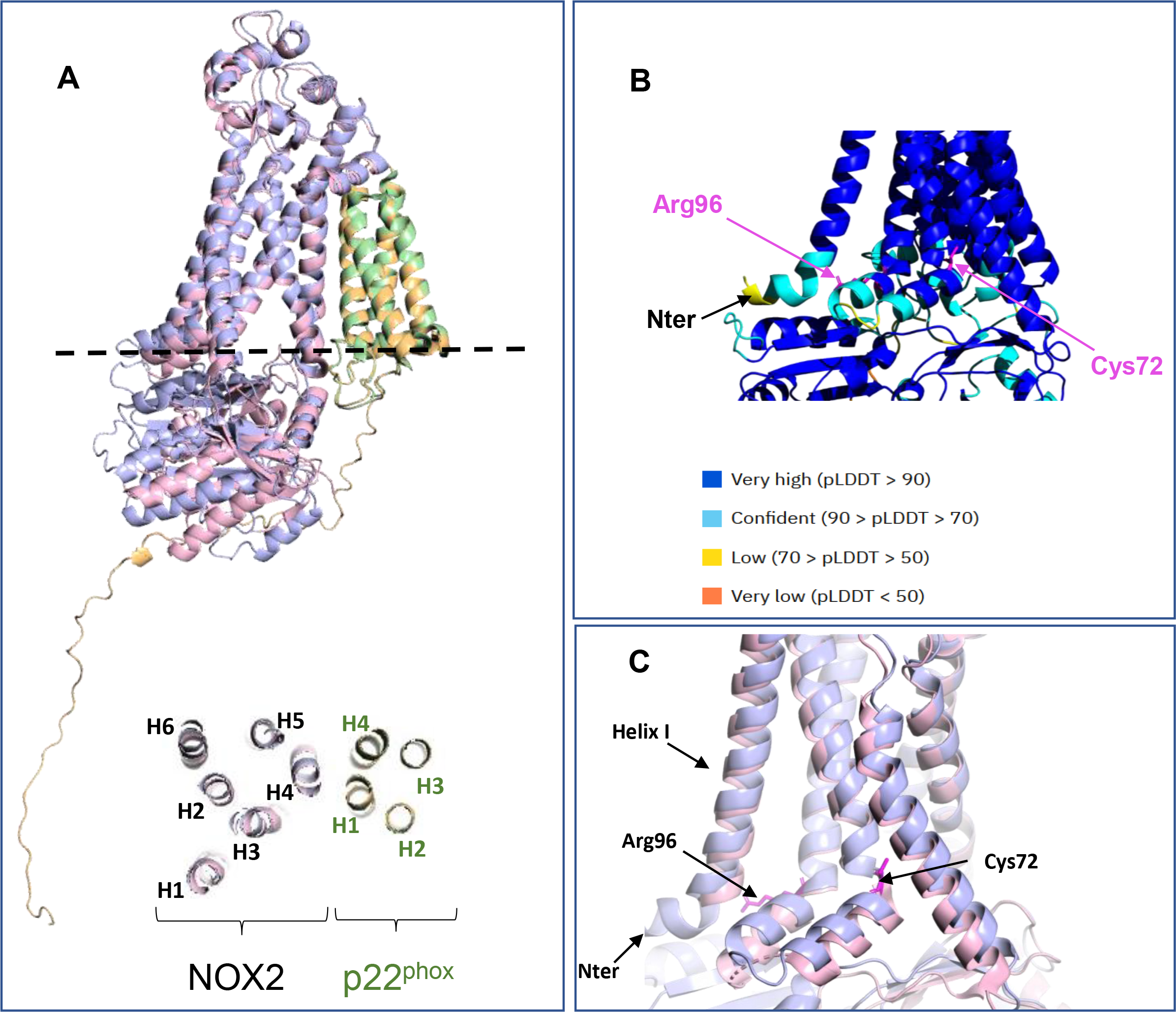
Comparison of the theoretical and experimental models of Cyt*b*_558_. **A. Superimposition of the selected model generated by locally-implemented AF2 and the experimental (8GZ3) cryo-EM structural model of Cyt*b*_558_.** Alignment was performed with PyMOL software based on the membrane region of NOX2. NOX2 and p22^phox^ from the selected AF-Cyt*b*_558_ model are colored in blue and orange, respectively. NOX2 and p22^phox^ from the experimental 8GZ3 model are colored in pink and green, respectively. *Bottom*: Superposition of the top view of the cross-section of AF and experimental models of Cyt*b*_558_ on the transmembrane layer at the position indicated as a dashed line (at the level of Trp106 from NOX2). B. The transmembrane domains of the selected AF2-Cyt*b*_558_ model. The color code of pLLDT scores show the confidence levels. **C. New local insights of the transmembrane region of NOX2 resulted from the AF model (light blue) compared to the experimental model 8GZ3 (light pink) NOX2**. The N-terminus of NOX2, unresolved in experimental models, ends in a small horizontal helix, pulling helix I outwards. The 72-96 region is folded back towards the membrane in the AF model compared to experimental model.

In the AF2 model, the predicted membrane domains showed a very high level of confidence (pLLDT>90) except for the cytosolic side at the interface between the transmembrane and DH domains with some lower pLLDT scores (70 <pLDDT< 90) (**Figure 2B**). These slightly lower confidence in the AF2 model match unsolved regions in the experimental structures. The N-terminus of helix I, for example, is unsolved in the cryo-EM structures, and was predicted by AF2 to terminate with a bend, such that the end of the helix is oriented parallel to the membrane stabilizing the C-terminal region of NOX2 (**Figure 2C**). Likewise, the region Cys72-Arg96 in NOX2 is organized in the AF2 model in two small hairpin helixes linked with the loop between Ser83-Ser87, also experimentally unsolved (**Figure 2C**).

The major differences between the AF2 model and the experimental structure of resting NOX2 (8GZ3) are located in its DH domain, which was unsolved in the 7U8G experimental structure due to its flexibility (**Figure 3A**). Between both models (AF2 and 8GZ3), the DH domains do not superimpose, due to a significant deviation in their orientation with regard to the membrane. The plasma membrane was predicted using PPM (23). Compared to the experimental structure of cyt*b*_558_ in the resting state, the DH domain of the AF2-cyt*b*_558_ was found closer to the transmembrane domains of Cyt*b*_558_. The above-mentioned and hairpin N-terminus regions were pushed up towards the membrane leading to a more compact form of AF2-cyt*b*_558_ (**Figure 3B**). DynDom (24) was used to calculate the deviation in the angle between the DH domain and the membrane between the resting (8GZ3) NOX2 and AF2 NOX2 models. This deviation angle was found to be of 17.6° (**Figure 3A**). Similar calculation between the orientation of the DH of the resting NOX2 (8GZ3) and of DUOX1 in the presence of calcium (36) (PDB code: 7D3F) lead to a deviation of 49.4°. DUOX1, one of the isoforms of NOX2, is considered to be in the active form in the presence of calcium, with the DH domain orientation corresponding to a signature of the activated enzyme. In the AF2-generated model, the DH domain adopts an intermediate position between the position of the DH domain of DUOX1 (active), and that of experimental NOX2 (inactive) suggesting that the AF2-generated model may not correspond to Cyt*b*_558_ in its resting state, but rather approximates the DH domain orientation of an intermediate between the fully active configuration and the inactive structure of NOX2.

**Figure 3.**
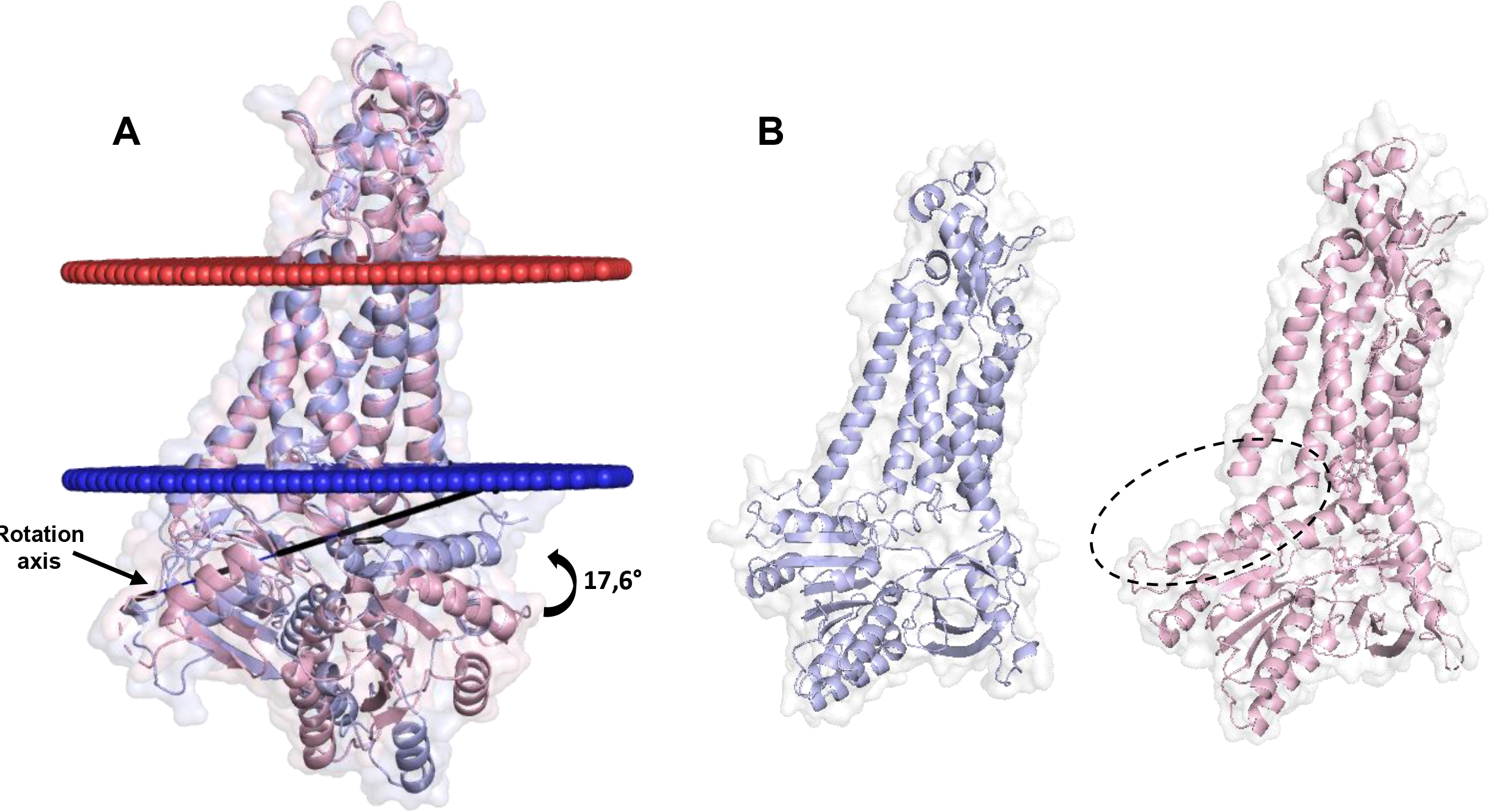
Focus on AF2 NOX2 model. **A. Comparison of the AF2-NOX2 (in blue) and experimental cryo-EM (8GZ3, in pink) models positioned into predicted membrane.** The rotational axis of the AlphaFold2 model of NOX2 relative to the experimental 8GZ3 structure was determined by using the DynDom server. In this analysis, the transmembrane domain is held as a fixed entity, while the tilt of the DH domain is measured at a displacement angle of 17.6 degrees. **B.** The AF2 model in blue is more compact than the experimental structure in light pink. The dashed region highlights the empty space between the dehydrogenase and the membrane domains correlated to the relaxed structure of NOX2 in the experimental structure.

### Insertion of the cofactors in the AF2-generated NOX2 model

The AlphaFill algorithm was used to integrate the cofactors (two hemes, one flavin and NADPH) into the AF2-NOX2. Alphafill considers only structural homolog with more than 25 % sequence identify. Therefore, at the time of the modelling, AlphaFill was unable to insert the hemes correctly. However, we expect this bottleneck to be solved when the AlphaFill database will be updated to include the two recent NOX2/p22^phox^ structures (5, 6). Due to the high superimposition of the transmembrane domains, both hemes are expected to adopt a position similar to what was observed in the two experimental structures. On another hand, by using Alphafill, flavin and NADPH could be inserted and optimized by using an Alphafill option based on energy minimization (**Figure 4A**). Flavin was added on the basis of the structure of the DH domain of NOX5 (PDB code: 5O0X)(38).The substrate NADPH was integrated on the basis of a flavin-containing protein (PDB code: 4YRY) (39). The optimization permitted to reduce the troubleshooting of structural clashes of the NADPH position but is not yet satisfactory. If AlphaFill is a powerful tool to insert cofactors in unsolved protein structure, the fact that it is based only on sequence identity remains a strong limitation.

**Figure 4.**
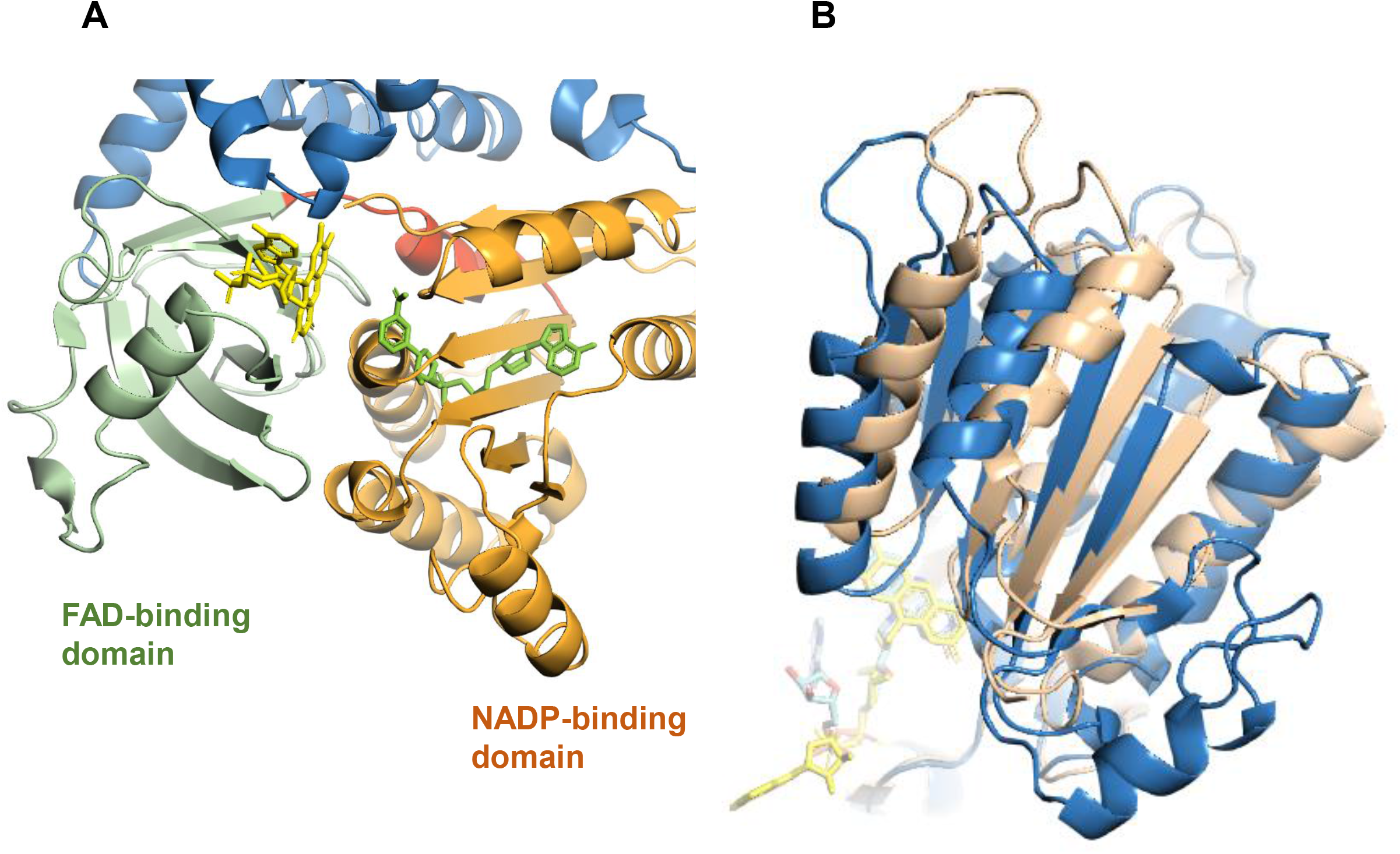
The DH domain of NOX2. **A. Predicted Flavin and NADPH binding positions into the AF2-generated NOX2 after energy minimization by AlphaFill.** The DH domain of NOX2 consists in two distinct domains, a FAD-binding domain (residues 295 and 390) coloured in pale green and a larger NADP-binding domain (residues 403-540) coloured in orange. Flavin and NADPH inserted using Alphafill are in yellow and green respectively (in blue are colored the membrane domains). Both domains display a typical dinucleotide-binding fold and are connected through a linker (residues 390-395 in red) that may act as the flexible hinge. In addition, there are regions interposed between the two domains lacking secondary structure elements (residues 342-350 and 314-324), suggesting a flexible interaction between the domains. **B. Superimposition of the DH domain based on the FAD-binding domain of the AF2 and 8GZ3 experimental models showing the deviation of the NADPH binding domain.** Predicted cofactors binding position into the AF NOX2 in comparison with the experimental 8GZ3 NOX2. The AF2 model is colored in blue with the flavin which is shown in yellow. The experimental structure is displayed in light brow with the flavin in light blue. When the FAD-binding domain of the DH domain of AF2 and experimental models are superimposed, the NADPH-binding domains were adopting a different orientation and deviated from about 9 Å

Despite this limitation, several interesting information could be drawn from the comparison of the DH domains between the AF2 model and the experimental structures. A structural superimposition of the theoretical and experimental resting NOX2 (8GZ3) based on the membrane domains highlighted the different position of FAD in both structural models mainly based on the above-mentioned DH deviation (**Figure S2**). Interestingly, the superimposition of the DH domain on the basis of the FAD binding site showed more distant FAD and NADPH pockets in the experimental resting NOX2 compared to AF2 model (**Figure 4B and 5A**). In the AF2-NOX2 model, the edge-to-edge distance between flavin N5 and NADPH could be estimated of about 7.1 Å (**Figure 5B**). No comparison is possible with the experimental structure of the resting Cyt*b*_558_ due to the absence of NADPH. In NOX5, this distance could be estimated by considering the C-terminus Trp695 proposed to mimic the nicotinamide NADPH binding (38). The distance between Trp695 and flavin N5 in the DH domain of NOX5 was found to be about 4 Å. In the experimental 3D structure of DUOX1, this distance was found to be about 10 Å, too far for hydride transfer. The distance between FAD and NADPH in AF2-cyt*b*_558_ is intermediate between NOX5 and DUOX1 but is still slightly too great to allow the expected efficient hydride transfer between NADPH and FAD. Unless the two pockets move even closer together on activation, it is interesting to note, in the AF2-NOX2, the presence of the C-terminal F570, unresolved in the experimental structure of resting NOX2 (**Figure 5B**). Interestingly, the orientation of the C-terminus of AF2-NOX2 brings Phe570 into close proximity to the flavin (4.5 Å) halfway between FAD and NADPH (**Figure 5B**). In this configuration, F570 stacks the isoalloxazine ring and could participate in the electron transfer regulation process between hydride transfer from NADPH and FAD.

**Figure 5.**
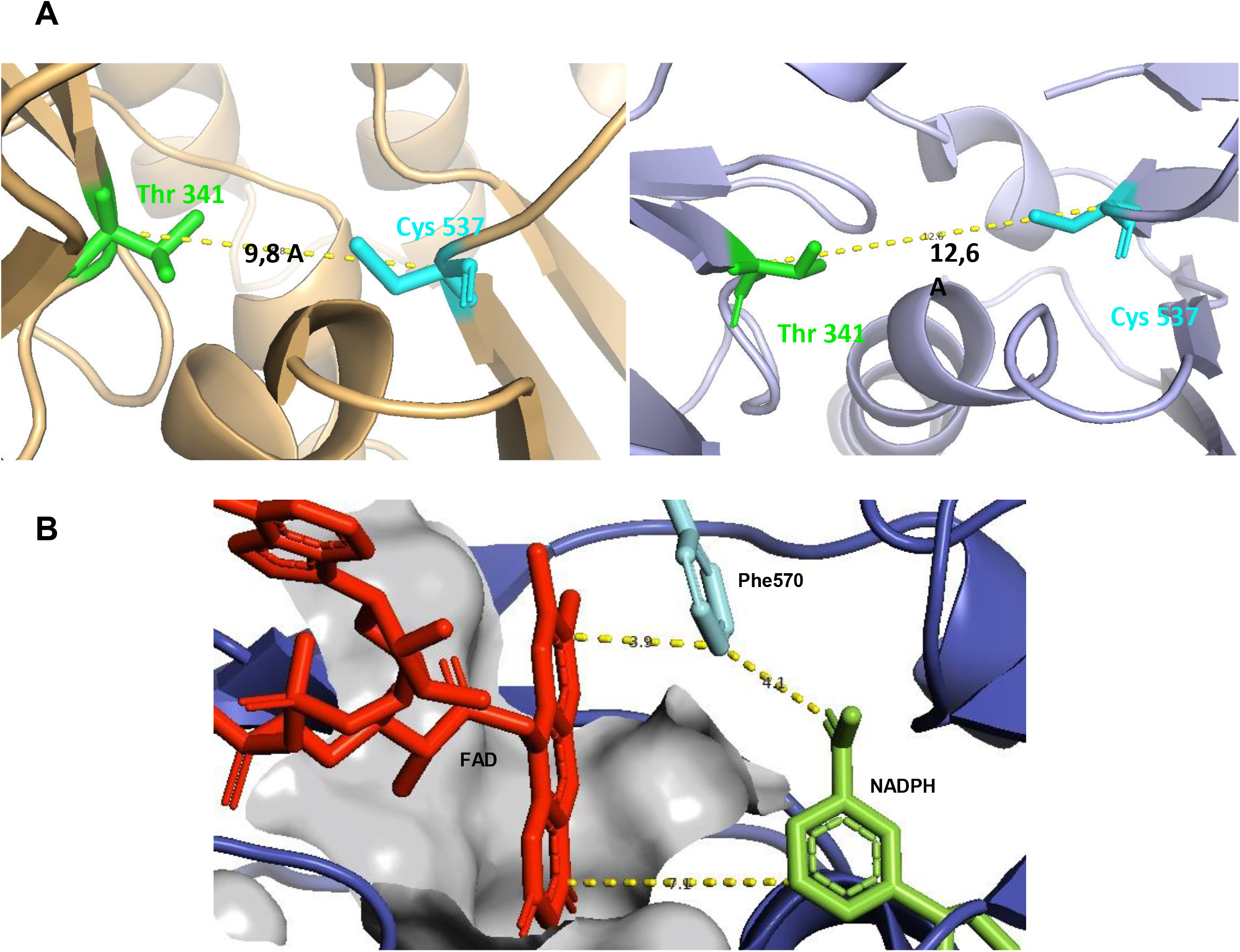
The FAD and NADPH binding pockets. **A. Superimposition of the FAD and NADPH binding pockets of the AF2 (left) and the 8GZ3 experimental (right) models.** The distances between FAD and NADP binding site were estimated on the distance between Cα of Thr341 belonging to the FAD-binding site and of Cys537 part of the NADP binding pocket. **B. Potential hydride transfer pathway from NADPH to FAD implying aromatic residue.** The FAD binding region from the AF (yellow) is colored in grey. The AF2-predicted NADPH is colored in green. The C-terminus Phe570 colored in light blue stacks the isoalloxazine ring of the flavin from the AF model at a distance of 3.9 Å.

### AF2-generated model of cytb_558_ with p47^phox^

Through regions involved in protein-protein interactions (such as its SH3 domains), domains interacting with lipids (such as its PX domain) and unstructured regions, p47^phox^ plays a central role in conducting the initial assembly and activation steps of the NADPH oxidase. AF2 was used to predict models of p47^phox^ in interaction with the complex formed by p22^phox^ and NOX2. Five different models of the multimer were generated, featuring a structurally stable transmembrane portion for NOX2 and p22^phox^ (as illustrated by the pLLDT scores), while the position of p47^phox^ varies from model to model (**Figure S3**). The DH domain showed the same orientation as the one observed for the AF2-Cyt*b*_558_ model suggesting that the docking of p47^phox^ did not impact the orientation of the DH domain. The selection of the most appropriate model was based on the criteria described in the literature for p47^phox^ and cyt*b*_558_ interacting regions, i.e. the presence of an interaction of the PRR region of p22^phox^ with the SH3 of p47^phox^, as well as a rotation of p47^phox^ orienting its PX domain towards the membrane. The models lacking these two interactions were not considered. The most appropriate model (Model 2 in **Figure S3**) displays interactions between the proline-rich region, located at residues 151-160 in the cytosolic C terminus of p22^phox^ and the SH3 domains of p47^phox^ as described extensively in the literature.

Spatial positions of cyt*b*_558_ with respect to the lipid bilayer were optimized by the PPM 2.0 method that accounts for the hydrophobic, hydrogen bonding and electrostatic interactions of the proteins with the anisotropic water-lipid environment described by the dielectric constant and hydrogen-bonding profiles. This simulation allowed a better visualization of the orientation of p47^phox^, in particular the PX domain, relative to the membrane (**Figure 6**). The AF2 model highlights the very high flexibility of p47^phox^. Once in interaction through its SH3 domains with the PRR region of p22^phox^ (including P152, P156 and R158 as shown in (40) (**Figure 7A**), p47^phox^ extended up to interacting at its C-terminal end with the 491-500 loop, and with the 451-458 loop of NOX2 and harpooning the membrane via its N-terminal PX domain.

**Figure 6.**
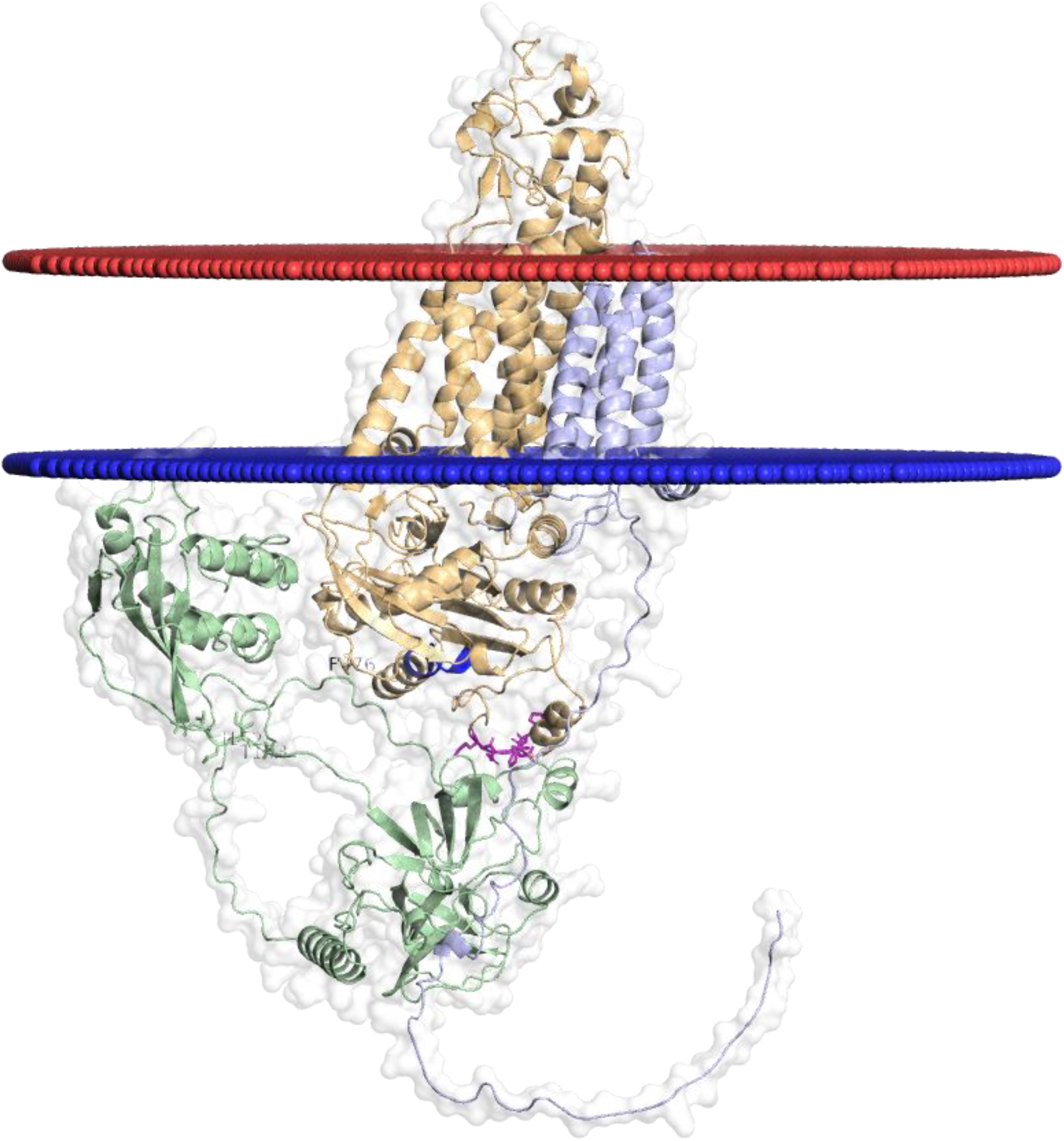
Selected AF2 model of the NOX2-p22^phox^-p47^phox^ complex. **A.** Global model of the interactions between NOX2 (light brown), p22^phox^ (light blue) and p47^phox^ (light green) inserted into a membrane predicted by PPM server. The pictures showing the 494-498 NOX2 region (purple) and the 451-458 NOX2 region (blue) of NOX2 interacting with p47^phox^ Cter region are enlarged in figure 7 (B and C, respectively).

**Figure 7.**
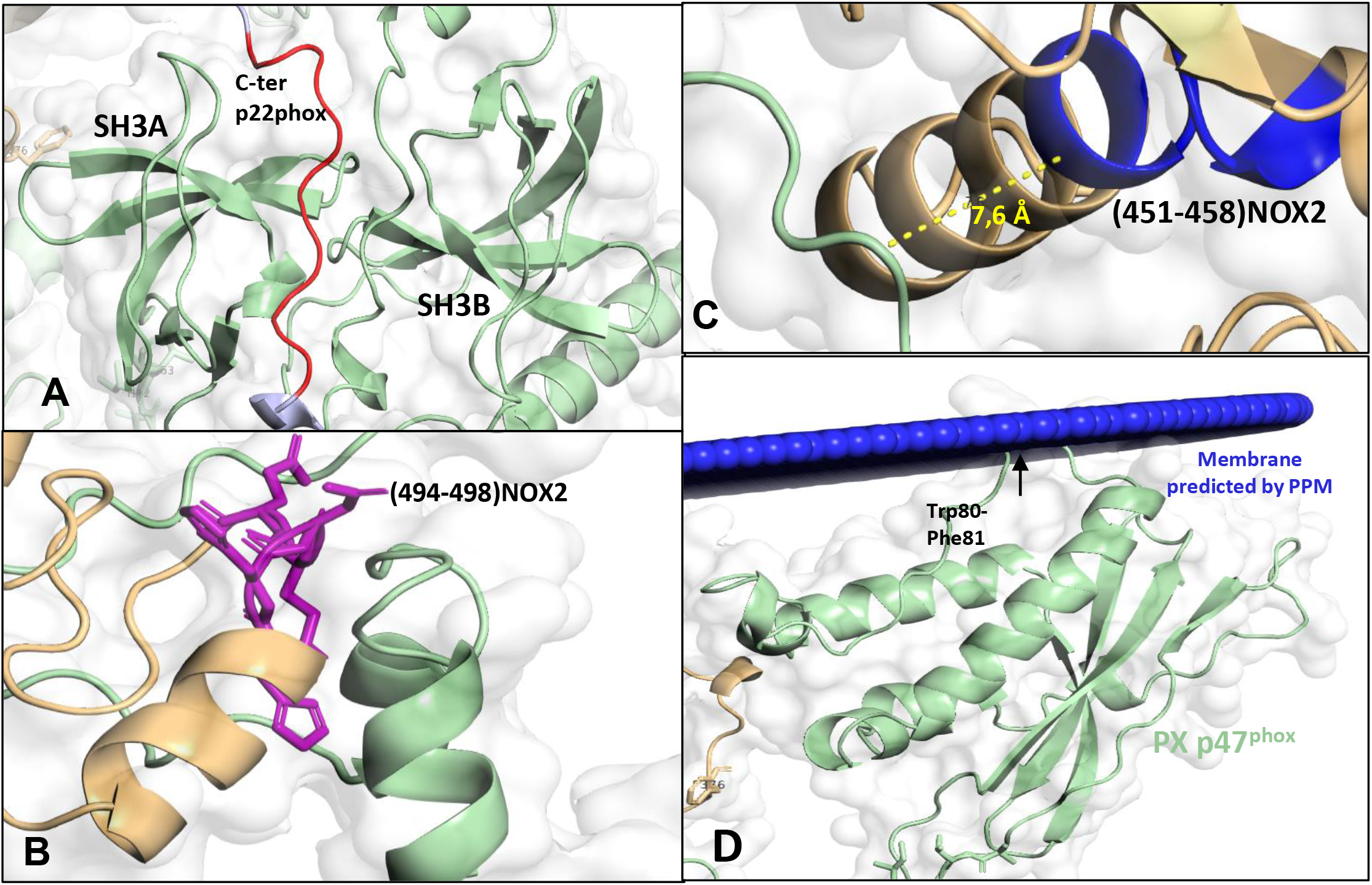
Interacting regions of p47^phox^ and cyt*b*_558_ in the AF2 model of NOX2-p22^phox^-p47^phox^ complex. **A.** Interaction of PRR region of p22^phox^ (red) and SH3 domains (green) from p47^phox^. **B.** Interaction between the 494-498 NOX2 region (purple) and the p47^phox^ Cter region (green). **C I**nteraction between 451-458 NOX2 region (blue) and p47^phox^ Cter region colored in green. **D.** Interaction of the K79-R85 region of the PX domain of p47^phox^ with the membrane via its hydrophobic W80-F82 and the contribution of K79 (green).

The above-mentioned interacting regions between p47^phox^ and NOX2 are in good agreement with the literature. The region corresponding to the 491-500 region (“HFAVHHDEEKD” residues) (41) is located in the DH domain of NOX2 (**Figure 7B**). It belongs to the NIS (*N*ADPH-domain *I*nsertion *S*equence) corresponding to 484–504 residues (42, 43), that is shifted of about 10 Å in the AF2 model of NOX2 compared to the experimental structure. The 491-500 region composed of charged amino acids, shows very close structural proximity to the C-terminal region of p47^phox^, with a distance of around 4 Å between the two proteins. The observed proximity suggests possible electrostatic interactions that supports the function of this sequence in appropriate NADPH oxidase assembly (44). The 451-458 region in the DH domain of NOX2 (45) located approximatively 7 Å away from to the C-terminal region of p47^phox^ may suggest a possible interaction with NOX2 (**Figure 7C**). These two regions 494-498 and 451-458 seem to play a role in the formation of the complex. However, these conclusions are based on a specific model which has been judged to be the most faithful in relation to the interactions already identified for this complex. Further studies involving other structures and experiments are required.

Experimentally, the 149-154 region (in particular Ile152 and Thr153) of p47^phox^ was proposed to play a crucial role in the production of superoxide anions by the phagocyte NADPH oxidase (46). However, in the AF2 model, this region was not found in close interaction with either p22^phox^ or NOX2 (**Figure S4A**). At least in the absence of other subunits, this region plays no role in the interaction between p22^phox^ and p47^phox^, nor in the process of translocation to the membrane. Its specific role remains unknown, but one hypothesis suggests that it may be involved in the interaction between NOX2 and p47^phox^ after the interaction between p22^phox^ and p47^phox^ has taken place (47).

The region 86-94 located in the B loop of the NOX2 protein was also proposed to interact with p47^phox^ (**Figure S4B**) (45). In the model predicted by AlphaFold2, this loop was found anchored in the membrane and has a large distance to p47^phox^. Another region of NOX2 proposed to interact with p47^phox^ is region 559-565 at the terminal region **(Figure S4B**). However, in the proposed AlphaFold2 model, this region was also distant from the p47^phox^ protein. These results do not support the involvement of these two regions in the NOX2/p47^phox^ interaction.

Finally, in the AF2 model, the PX domain is anchored in the membrane not via the I65 residue as expected (48), but via a hydrophobic region (W80-F81) and a positively charged residue (K79) (**Figures 7D and S5**). This anchoring may contribute to bringing the critical residues involved in canonical recognition of phosphoinositides (PtdIns3P) “RPPR” (R43, P73, P76, R90) and those binding potentially non-canonical phosphoinositides (H51, K52, K55, R70).

### MD simulations of the NOX2-p22^phox^-p47^phox^ complex embedded into a membrane bilayer

This predictive AF2 model emphasizes that AlphaFold2 implicitly considers the membrane and the hydrophobic interactions that membrane and cytosolic proteins can develop with their environment during the assembly process. However, considering that some of our results are in contradiction with the experimental literature and the fact that AF2 only produces static models, we proceeded to molecular dynamics (MD) simulations to observe the dynamics of the structure more closely and to explore the lipid-protein interactions, to enhance our understanding of the complex behavior within the membrane environment. AF2 predictions suggested that the anchorage of the PX domain does not take place via the I65 residue as expected (48) but via the K79-R85 region of the p47^phox^-PX domain, in particular the hydrophobic (W80-F81) and the positively charged (K79) residues (**Figure 8** and **Table 2**).

**Figure 8.**
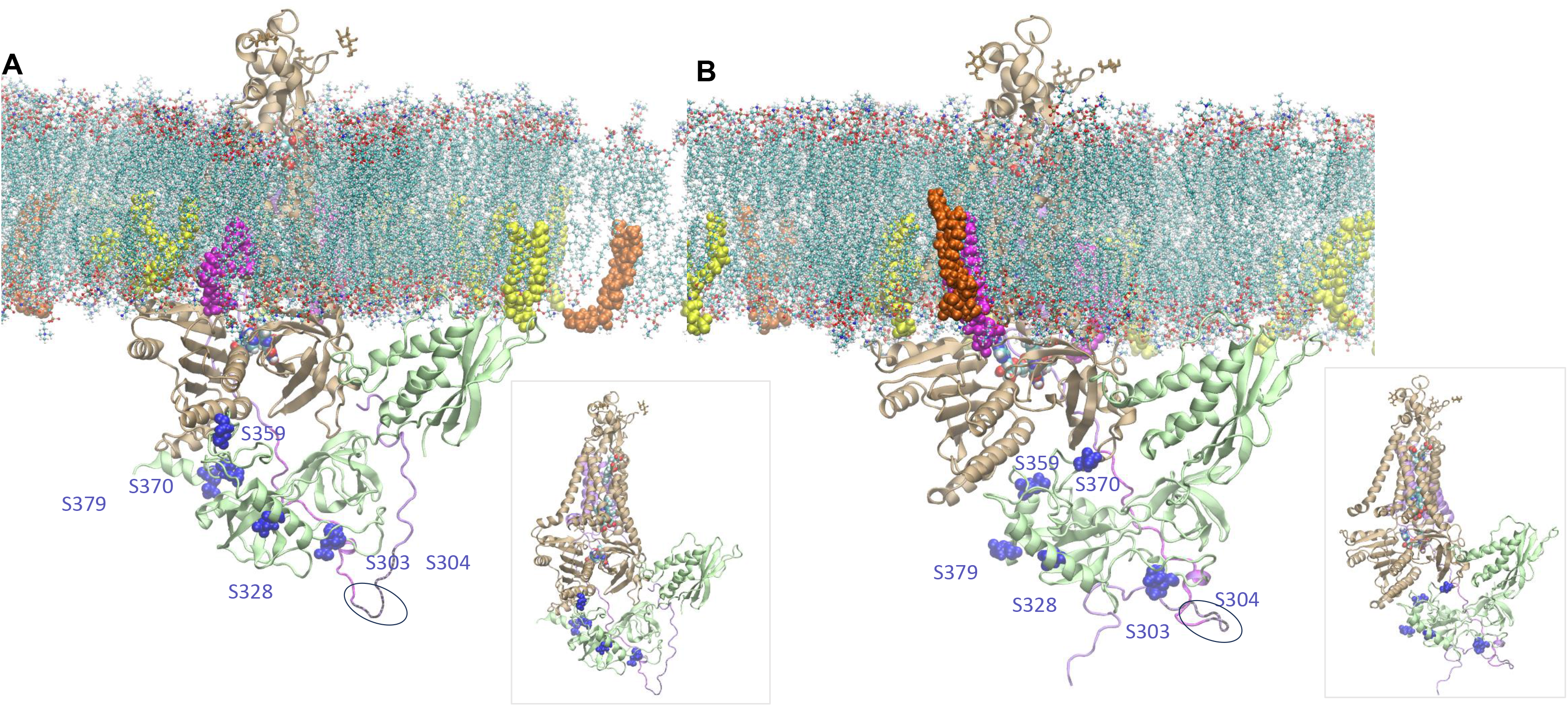
The p47^phox^-PX domain interactions with the lipid bilayer. **(A) A snapshot from NOX2-p22^phox^-p47^phox^ MD simulations replica 1, (B) A snapshot from NOX2-p22^phox^-p47^phox^ MD simulations replica 2.** The inset figures show the same conformation without the membrane components for clarity. NOX2 protein is colored in light brown, p22^phox^ in light blue, and p47^phox^ in light green, all depicted in cartoon. The POPS, PI(3)P, and PI(3,4)P2 lipids are shown in spheres, and are colored in yellow, magenta, and orange, respectively. The phosphorylated p47^phox^ residues are shown as purple spheres. The glycans N132, N149, and N240 of NOX2 protein are colored in gold and shown as sticks. The p22^phox^ K149-E169 region is colored in magenta. The p22^phox^ E169-G177 region is encircled and colored in gray. Hemes and FAD are colored according to atom name and are shown as spheres.

Throughout the 500 ns MD simulations, the critical residues involved in the canonical recognition of phosphoinositides, PXXP (P73-P76) and the K79-R85 region of the p47^phox^-PX domain, as well as F18-Q22 of PX domain and P128-K134 of the linker connecting PX and SH3, interacted with the bilayer. Since the position of the specific lipids (PI(3)P, PI(3,4)P2, POPS etc.) fluctuated significantly in the lipid bilayer, we have not sampled the particular interaction of these canonical residues and the PIs due to the random positioning at the beginning of the simulations.

We observed persistent anchoring of the hydrophobic W80-F81 residues of the p47^phox^-PX domain within the membrane bilayer of the NOX2-p22^phox^-p47^phox^ complex in our MD simulations. Also, I65 got closer to the bilayer, interacting with NOX2 through the part connecting the lipid embedded TM6 and the FAD domain (through V296, K313, K315).

The persistent interactions observed, particularly between the p22phox PRR and the SH3 domains of p47phox, as well as the interaction of p47phox with the 491-500 loop of NOX2, provide support to the structural models and highlight the importance of these regions in maintaining the structural integrity and activation of the NADPH oxidase complex. We did not observe a stable interaction between p47^phox^ C-terminal and the 451-458 loop of NOX2 in either of the simulation replicas. However, in one instance, these regions approached close proximity, suggesting the potential for transient or weak interaction under certain conditions.

The interaction dynamics of the critical regions in the phosphorylated state, particularly a lack of interaction between the p47phox region 149-154 and the NOX2 region 377-379 and the interaction of p47^phox^ PRR (K360-P369) with the NOX2 E375-P383 region may facilitate the recruitment or positioning of p47^phox^ for subsequent binding with p67phox, that may lead to the complex’s activation. Future simulations, including the unphosphorylated state starting from the same structural model, might shed further light on these intricate processes.

These interactions, while corroborating the AF2-generated structural predictions, also highlight the dynamic nature of the NOX2-p22^phox^-p47^phox^ complex and its regulation by phosphorylation states. Our findings emphasize the importance of considering both static models and dynamic simulations to fully understand the molecular mechanisms governing the assembly and activity of the NADPH oxidase complex.

### AF2-generated model of cytb_558_ with p47^phox^, p67^phox^ and Rac1

We attempted to predict the assembly of the p47^phox^, p67^phox^ and Rac proteins with cyt*b*_558_ but unfortunately, this was unsuccessful, perhaps due to the large number of flexible and unstructured regions present in these proteins. The models obtained proved unsatisfactory as they took the form of protein “spaghetti”, with low levels of PLDDT scores. Therefore, we opted for an alternative way of obtaining a model of the complete complex in its active form, bypassing the number of unstructured regions and thus facilitating AlphaFold2 modelling.

To do this, we used the chimeric protein called Trimera (**Figure 9A**). By combining the essential functional domains of p47^phox^, p67^phox^ and Rac1Q61L, Trimera is well known and canonically used to activate the complex experimentally in vitro, although it requires an additional activator, the arachidonic acid (17). Thus, we hypothesized that using Trimera in AF2 modelling may provide a simplified but representative model of the active NOX2 enzyme complex.

**Figure 9.**
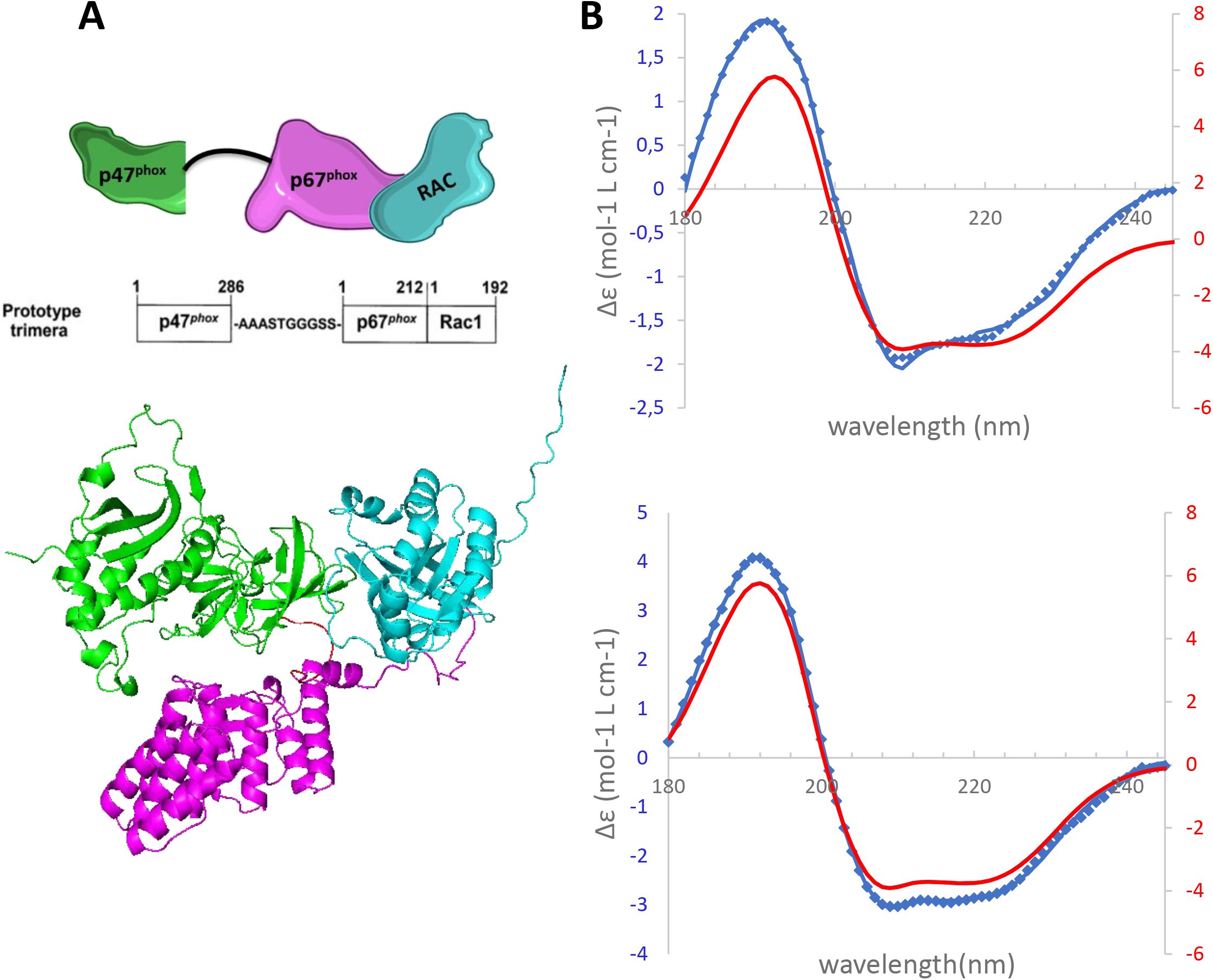
Conformation studies of Trimera. A. Scheme and AF-generated model of the Trimera. The trimera was designed to gather the essential domains of the cytosolic proteins which consist in the PX domain and the two SH3 of p47^phox^ (in green from residue 1 to 286), linked by a linker to the TPR motifs and the AD region of p67^phox^ (in magenta from residues 1 to 212). The latter is linked directly with the full-length Rac1 protein (colored cyan) which contains the Q61L mutation. **B. Experimental and theoretical CD spectra of Trimera.** The theoretical spectrum of the AF model of Trimera calculated using SESCA software was colored in red. The experimental SRCD spectra of Trimera in the absence (A) and the presence of arachidonic acid (B; 400 µM) was colored in blue.

The online tool AlphaFold2 CoLab was used to predict the spatial organization of the Trimera alone in order to compare it to experimental conformational analysis using SRCD approach. The five proposed models show a globally consistent superposition with a preserved structure of structured domains, but their orientation differs from model to model depending on the positioning of linkers and unstructured regions (**Figure S6**). Selection of the optimal model was based on an evaluation of several key criteria, aimed at ensuring structural compatibility with known structural features (**Figure 9A**): (i) same-side orientation of the PB region of Rac1 and the PX domain of p47^phox^ for membrane interaction; (ii) accessibility of the SH3 domains of p47phox and the AD region of the p67^phox^ protein for interaction with the membrane proteins. CD analysis was performed on purified Trimera in the presence or absence of arachidonic acid. As shown in **Figure 9B**, the theoretical and experimental CD spectra obtained in the presence of arachidonic acid show similar characteristics, suggesting that the conformation of the optimal model of Trimera generated by AF2 should adopt an active conformation.

Following this encouraging result, the NOX2-p22^phox^-Trimera multimer was modelled using AF2. The five models proposed for this interaction are very similar and superimposable, with very satisfactory PLDDT scores. Above all, the key interactions between membrane proteins and cytosolic proteins described as important for activation of the NOX2 enzyme complex in the literature are present (**Figure 10A**): (i) the interaction of the C-terminal PRR region of p22^phox^ with the SH3 domains of p47^phox^, important for the translocation of p47^phox^ and p67^phox^, is present and is high lightened in black in **Figure 10A** ; (ii) the activation domain (AD), corresponding to the “VAQLAKKDYLGKATVVASVVD” sequence of p67^phox^ (colored in magenta in **Figure 10A**) is interacting with the region (residues 357-371, colored in light brown) of the NOX2 DH domain. This interacting region was described as essential for NOX2 activation (49); (iii) the PX domain of p47^phox^ is positioned on the membrane side with the same interaction with the lipid bilayer as observed in the NOX2/ p22^phox^/p47^phox^ model.

**Figure 10.**
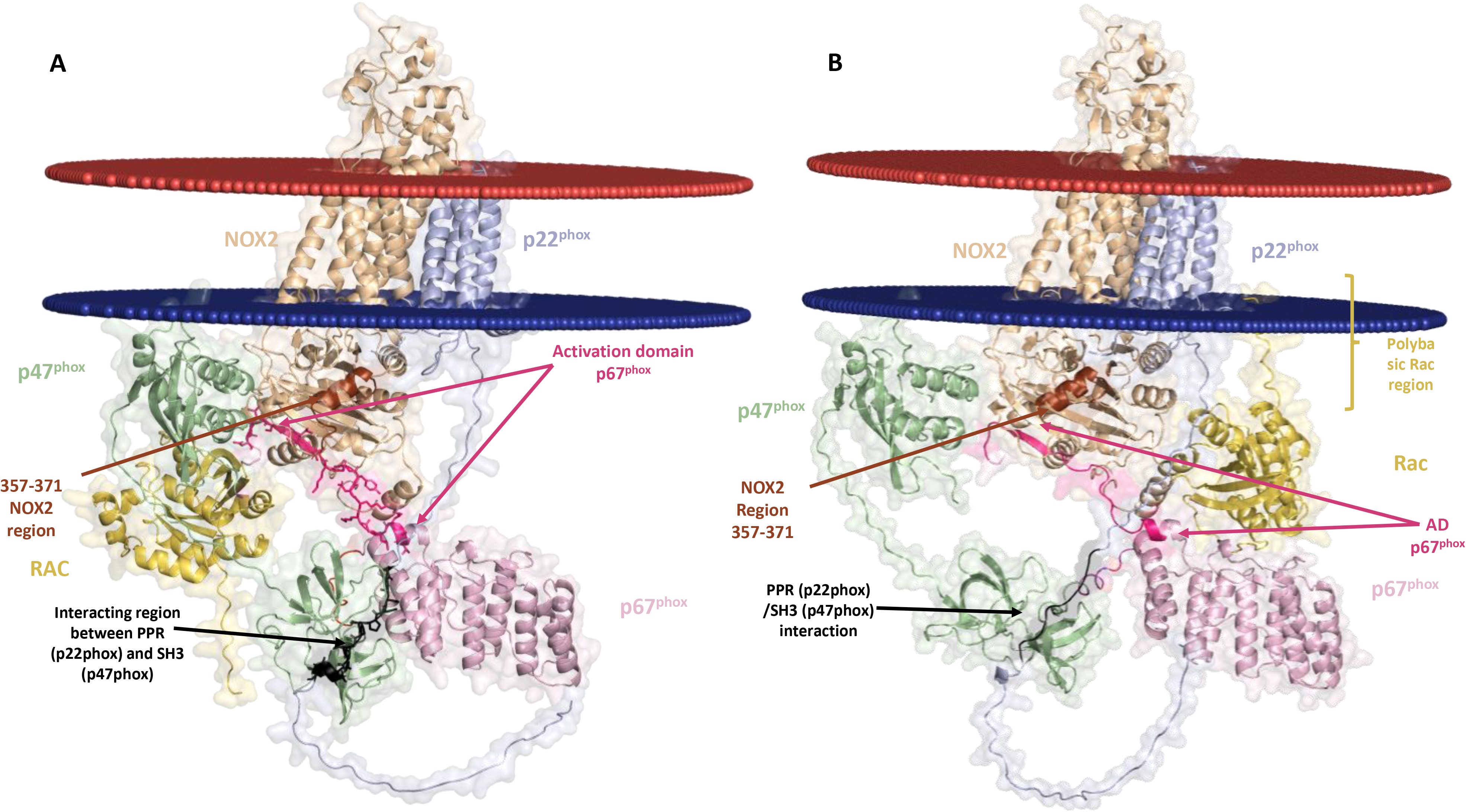
A. AF-generated model of NOX2, p22^phox^ and Trimera multimer inserted into a predicted membrane. Full model of the interaction between NOX2 (light brown), p22^phox^ (light blue) and Trimera (fusion of p47^phox^ in light green, p67^phox^ in light pink and Rac1 in yellow) inserted into a membrane predicted by PPM server. The p67^phox^ activation domain, the helix (357–371) of NOX2 and the PPR region of p22^phox^ were highlighted in pink, brown and black respectively. **B. AF-generated model of NOX2, p22^phox^, p47^phox^/p67^phox^ dimer and Rac1 multimer inserted into a predicted membrane.** The full model consists in the NOX2 protein (light brown), the p22^phox^ protein (light blue) and fused essential domains of p47^phox^ (1-586, colored in light green) and p67^phox^ (1-212 colored in light pink) and the separated Rac1 protein (yellow).

However, the positioning of Rac is not optimal. In this configuration, its position does not allow the expected interaction of its polybasic region with the membrane. This suggests that in the complex formed with the trimer, it is not possible for the AD region of p67^phox^ to interact with NOX2 at the same time as Rac interacts with the membrane. This may be explained by the too short distance between Rac and p67phox to give them sufficient flexibility to move into the right position. To circumvent this limitation using AlphaFold2, an alternative model of the complex was explored by splitting Trimera between Rac and the essential p47^phox^/p67^phox^ regions (**Figure 10B**). The C-terminal PRR region of the p22^phox^ protein (in black) still establishes a specific interaction with the SH3 domains of the p47^phox^ protein, a crucial interaction for orchestrating the first docking steps. Simultaneously, the activator domain (AD) of the p67^phox^ protein engages in a specific interaction with the 357-371 region in the DH of NOX2. This interaction plays a fundamental role in enzyme activation. Most importantly, the PX domain of the p47^phox^ protein and the polybasic region of the Rac protein both face the membrane. This configuration enables correct engagement with membrane phospholipids, which plays a central role in the coordinated interactions and activation of the complex. During the final drafting phase of this work, the 8WEJ structure of the activated complex was published in the PDB (https://doi.org/10.2210/pdb8WEJ/pdb). Although this structure lacks p47^phox^, it has several similarities with the AF2 model obtained and in particular a Rac/p67^phox^ interface very similar to that illustrated in figure 10b.

## Discussion

Although the crucial roles of p47^phox^, p67^phox^ and Rac1 or Rac2 in the cyt*b*_558_ activation process have been extensively detailed in the literature, the mechanism of cyt*b*_558_ activation is still a matter of debate with many unknowns. The respective involvements of the various subunits in enzyme activation is manifested by a series of complex protein-protein interactions: more specifically, the interaction between p67^phox^ and NOX2, as well as the interaction between p47^phox^ and p22^phox^ and Rac with NOX2 (2, 50, 51). Furthermore, lipid-protein interactions are essential, notably those of Rac and p47^phox^ with the membrane (13, 52). These interactions are complex and need to be synchronized with NOX2/p22^phox^ membrane proteins for optimal enzyme activity. By combining computational approaches and experimental data, we built the macromolecular edifice step by step, with the ultimate aim of obtaining a model of the assembled complex. Therefore, we used AF2 online and locally implemented to improve the models of complexes.

It is essential to stress that the AlphaFold algorithm still encounters difficulties when modeling proteins with high flexibility and numerous unstructured regions, as is the case for p47^phox^, p67^phox^ or the C-terminal region of p22^phox^(53). In this situation, literature data and existing structural information, even if fragmentary, become crucial assets for selecting the best model from among those proposed by the software, and for guaranteeing the quality and accuracy of the chosen model.

### Orientation of the DH domain and of the FAD-and NADPH-binding domains: hallmarks of active structures?

The DH domain of NOX2, homologous to the protein ferredoxin-NADP+ reductase, is made up of two distinct structured domains, a FAD-binding domain (residues 295-390) and a larger NADP-binding domain (residues 403-590). Both domains display a typical dinucleotide-binding fold and are connected through a linker (residues 390-395) that may act as a flexible hinge (**Figure 4A**). Two regions that are interposed between the two domains lack secondary structure elements (residues 342-350 and 314-324), thus suggesting a flexible link between the domains. Using AF2, we observed a significant deviation of the DH domain between the experimental “inactive” and AlphaFold2 models of NOX2, as well as a shortening of the distance between the flavin-binding region and the NADPH-binding region in the AlphaFold2 model. Superimposing the AF2 and experimental NOX2 models and DUOX1 structures reveals that in the predicted model, the DH domain adopts an intermediate orientation between the active form of DUOX1 and the inactive form of NOX2. This raises questions about the NOX2 state generated by AF2 since it could represent an intermediate form of NOX2, located between the active and inactive forms of the protein. However, it is also possible that the active form of NOX2 is different from that of DUOX1, since the mechanisms of calcium activation versus regulatory protein interaction (p67^phox^, p47^phox^, Rac) are different in both systems. In none of the complex models obtained, the positioning of the DH domain was affected by the presence of the other subunits compared with its positioning in Cyt*b*_558_ alone. However, the relative orientation of the DH domain to the membrane was stressed to be important for the activity of the NOX. This may support the hypothesis that AF2 generated an “active” form of Cyt*b*_558_.

To switch the DH domain of the experimental “inactive form” Cyt*b*_558_ to the potentially “active” AF2 form, two distinct arrangements of and within the DH domain might correspond to mechanistic steps of the Cyt*b*_558_ activation. One movement involves the entire DH domain, resulting in a repositioning of the DH closer to the membrane, while another involves bringing the FAD-and NADPH-binding lobes closer together to facilitate the hydride transfer from NADPH to FAD. It is interesting to note that, in the state of Cyt*b*_558_ predicted by AF2, the C-terminal arm of NOX2 shields the isoalloxazine ring from bulk solvent as the aromatic side chain of Phe570 faces the pyrazine ring of FAD (parallel π-stacking at a distance of 3.9 Å) at the potential nicotinamide binding site. The C-terminal part of NOX2, through the interaction of its C-terminal Phenylalanine (F570) with the isoalloxazine ring (also seen in the simulations), could contribute to a modulation of substrate binding and a control of the distance between FAD and NADPH to allow efficient hydride transfer. Indeed a certain flexibility of the C-terminus region of and within the DH domain, and a conserved aromatic residue at or near the C-terminus among members of the ferredoxin-NADP reductase (FNR)-like proteins were proposed to participate in the control of the NADPH binding and enzyme activities (54–56). This is also consistent with similar hypotheses described for the prokaryotic NADPH oxidase SpNOX (57). However, in NOX2, the sole experimental *in cellulo* data did not support this idea since it was shown that substitution of F570 did not significantly affect the NADPH oxidase activity (58). However more attention should be probably paid to the DH internal motion and the C-terminus end of NOX2. The C-terminal region of NOX4 has been shown to be one of the keys to explain the constitutive activity of NOX4 (59) and among the structurally related NOX homologs (NOX1-5), only NOX4, the sole constitutively active NOX enzyme, has a serine as a C-terminal residue while the others, which all require an activation step, have a phenylalanine (**Figure S7**). More extended studies are needed to solve this question.

### Exploring the interactions of the p47^phox^-PX domain with the membrane bilayer

The PX domains are known to target membranes through specific interactions with phosphoinositides. Its name derived from the phagocyte NADPH where it was originally identified (*phox*) (60). p47^phox^-PX domain has been shown to bind for a variety of phosphoinositides and lipids (61, 62). In p47^phox^-PX domain, NMR structure revealed the existence of an 8-Å deep cavity proposed to be involved in interactions with negatively charged phospholipids (63), such as PI(3,4)P2 and phosphatidic acid (64). Studies based on directed mutations and calculations of electrostatic potentials determined the key amino acids for the interaction of the PX domain with the membrane. Residues R43 and R90 were identified as essential for PX interaction with phosphoinositides while residue I65, adjacent to canonical and secondary phosphoinositide binding sites, enables non-specific penetration of neutral lipid monolayers (48). Here we showed that alternative interactions may occur between p47^phox^ and the lipid bilayer coupling specific interactions of the headgroup of PI3P with the region K79-R85 and the hydrophobic lipid insertion through W80-F81. The two regions of the p47^phox^ PX-domain involved in the membrane interaction might play a role in either protein translocation to the membrane or in complex stability. The previous report showing that mutation of R42 residues or mutations in the PXXP region do not abolish membrane association of p47^phox^ supports the idea of other alternatives for p47^phox^ to bind with the membrane (62). Due to the dynamic nature of specific lipids in the bilayer, MD simulations did not capture stable interactions with canonical residues; however, we consistently observed the hydrophobic W80-F81 residues of the p47phox-PX domain anchored within lipid bilayer. Although the associated functional role is not clear, we could believe that one event may play a weak allosteric role in p47^phox^ membrane binding and potentially also the orientation of the PX domain while the other one could contribute to overall membrane avidity and complex stability.

### Development of a 3D structural model for the complete NADPH oxidase complex

Using AlphaFold2, we produced several structural models of the different subunits of the complex, as well as models of the interactions between these subunits and implicitly protein-membrane interactions. Based on known interactions, such as the PRR of p22^phox^ with the SH3 of p47^phox^ and the AD region of p67^phox^ with the dehydrogenase domain of NOX2, we were able to validate or invalidate these models. In addition, by positioning a membrane model, we were able to verify the orientation of p47^phox^ PX and Rac polybasic domains in relation to the membrane.

In the AF2 model of the NOX2/p22^phox^/p47^phox^ complex, the p47^phox^ protein seems to develop few interactions with NOX2. However, based on biochemical studies, such as cell-free interactions, mainly four regions of NOX2 have been proposed to interact with the p47^phox^ protein. The residues 494-498 and 451-458 of NOX2 are found in the selected AF2 model to be interacting with p47^phox^ suggesting that they could play a role in the constitution of the complex. These interactions are seen partly in the simulations, but not in the AF2 model similar to the new active structure 8WEJ. One might suggest that after constitution of the NOX2-p22-p47 complex, these parts become available for p67^phox^ Activation Domain binding. However, these conclusions are based on a specific model that has been judged to be the most faithful in relation to the interactions already identified for the active complex. On another hand, the residues 86-94 and residues 559-565 of NOX2, mentioned in the literature as areas of interaction with p47^phox^, do not appear to be involved in the AF2 models and simulations as they are too distant from each other.

On the other hand, the regions of NOX2 that are spatially closer to p47^phox^ in our AlphaFold2 models, namely residues 494-498 and residues 451-458, could constitute potential areas of interaction. This result provides a promising starting point for future experimental studies aimed at gaining a better understanding of this hitherto little-studied interaction.

Interestingly, the model of cyt*b*_558_ interacting with the Trimera have shown that it is impossible to have simultaneous interaction of the activator domain (AD) of p67^phox^ with NOX2 and interaction of the polybasic region (PB) of Rac with the membrane. These spatial constraints may explain why the addition of arachidonic acid is necessary in the cell-free system to obtain an active complex (65), whereas Trimera can bind to simple membrane vesicles without membrane proteins (66). By replacing Trimera with a p47^phox^/p67^phox^ dimer and Rac separately in the AF2 modeling, we obtained a better model with Rac interacting with the membrane by anchoring its C-terminus into the lipid bilayer. We consider this model to be the most appropriate for generating a correct structure of the complex in its active form, as close as possible to the native enzyme.

The model presented here could provide the first structural description of the full assembled NADPH oxidase complex. This important step in the characterization of the complex provides important information regarding its fundamental assembly mechanisms, but also raises other questions. If the first steps are related to assembly processes, other organizational steps are probably necessary to obtain a functional structural organization. Further deep analysis of these models is required to assess their correlation with the many mutations leading to CGD, and also to identify the relevance of the various targets of the enzyme inhibitors.

#### Abbreviations

AA: *cis* arachidonic acid
AF2: AlphaFold2
AD region: Activation Domain region
AI: artificial intelligence
Cyt*b_558_*: cytochrome *b_558_*
PRR: proline rich region
SH3 domain: sarc homology 3 domain
PB: polybasic region of Rac
PL: phospholipid
PX: phagocyte oxidase domain
TPR: tetratricopeptide repeat
PBS: phosphate buffered saline
CGD: chronic granulomatous disease
SRCD: synchrotron radiation circular dichroism
*phox*: phagocyte oxidase
NADPH: nicotinamide adenine dinucleotide phosphate reduced
FAD: flavin adenine dinucleotide
SDS: sodium dodecyl sulfate
aa: amino-acid
ROS: reactive oxygen species

## Acknowledgments

We thank Dr. E. Pick for providing with the Trimera plasmid. We acknowledge the financial support of the ANR French agency (N°ANR-21-CE29-0013). We are indebted to the «Initiative d’Excellence» program (LabEx) PALM for SA’s PhD financial support (contract n°ANR-11-IDEX-003-02). SRCD measurements on DISCO beamline at the SOLEIL synchrotron facility were performed under proposal N°20211318. We acknowledge Frank Wien assistance from SOLEIL beamline staff for his help. This work was supported by the«Initiative d’Excellence» program from the French State (Grant «DYNAMO», ANR-11-LABX-011-01). BAF received funding from the European Union’s Horizon 2020 research and innovation programme under the Marie Skłodowska-Curie grant agreement No. 101034407 (COFUND FP-DYNAMO-PARIS). Part of the molecular dynamics simulations has been performed at TGCC-GENCI machine IRENE on grant no. A0150714651.

