## Supplemental file for "Combining computational biology and experimental knowledge to draw a structural profile of the active membrane-assembled NADPH oxidase complex"

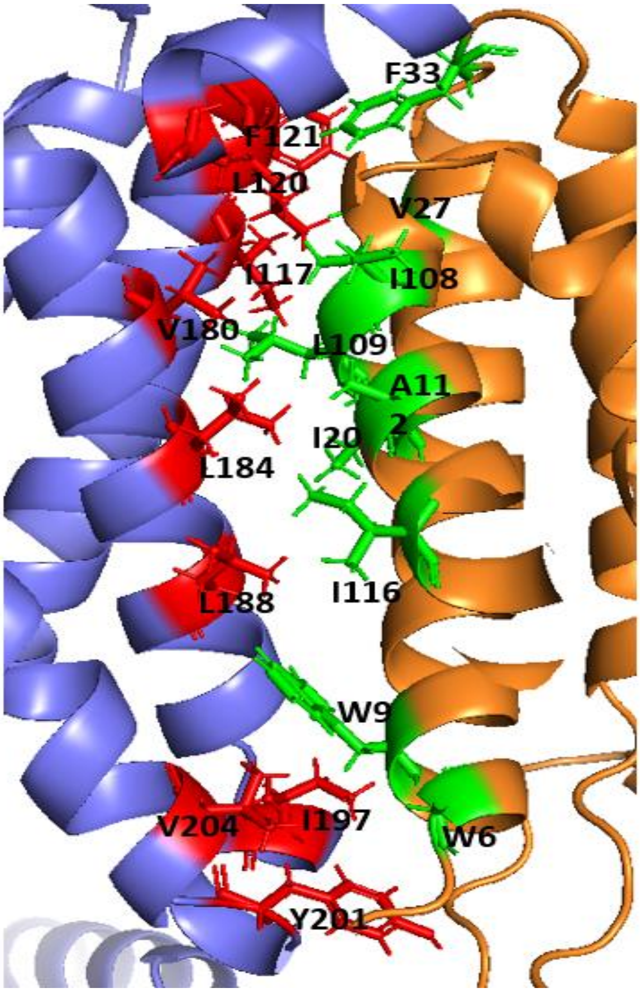


**Figure S1: A. Interacting region between NOX2 and p22$^{phox}$ in the AF2 model.** NOX2 is shown in blue and p22$^{phox}$ in orange, both illustrated in cartoon style. NOX2 residues involved in this interaction are highlighted in red, as sticks, while p22$^{phox}$ residues involved in this interaction are shown in green.

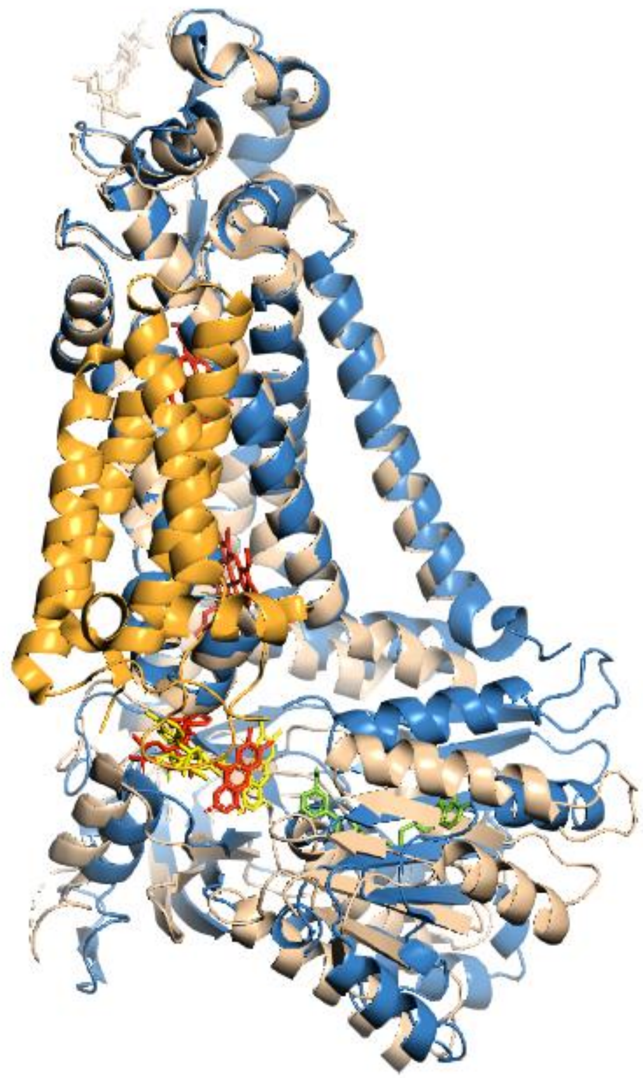

**Figure S2: Predicted cofactors binding position into the AF2 NOX2 in comparison with the experimental 8GZ3 NOX2.** The superimposition is done on the membrane domains. The AF2 model is colored in blue with the flavin and NADPH positioned after energy minimization by AlphaFill are shown in yellow and green, respectively. The experimental structure is displayed in light brown with the flavin colored in red. $p22^{phox}$ appears in orange.

.

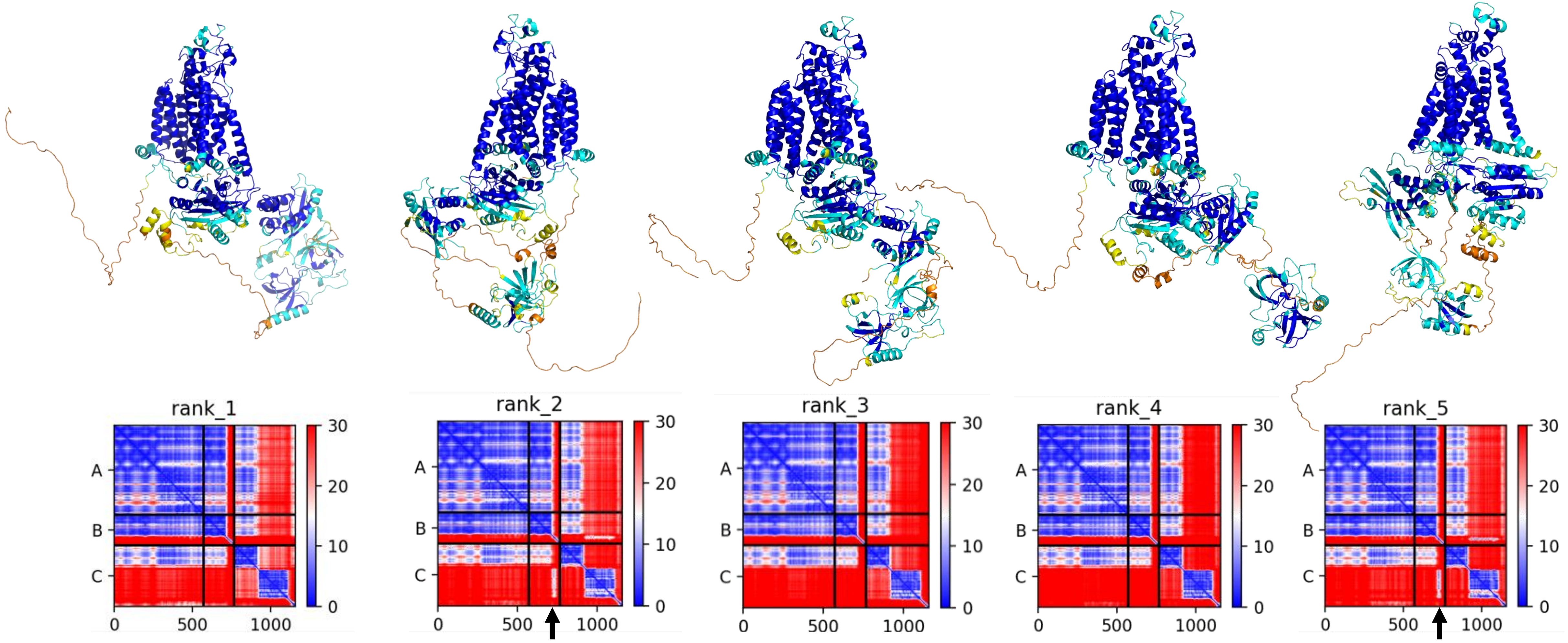


**Figure S3. AF2 predicted models (five) of the complex formed by p22$^{phox}$ associated with NOX2 and p47$^{phox}$.** The colors in the structural models describes the pLLDT score, showing the confidence levels of the different parts of the structure (same color code as in Figure 1) . **E**ach AF2 model is associated to a square that describes the expected errors in the distance of two residues in a 2-dimensional (pairwise) data set. Model # 2 (superimposable with model #5) was selected based on the criteria described in the literature for p47$^{phox}$ and cyt*b*$_{558}$ interacting regions. The arrows indicates the interacting region between the PRR region of p22$^{phox}$ and the SH3 of p47$^{phox}$.

**A**

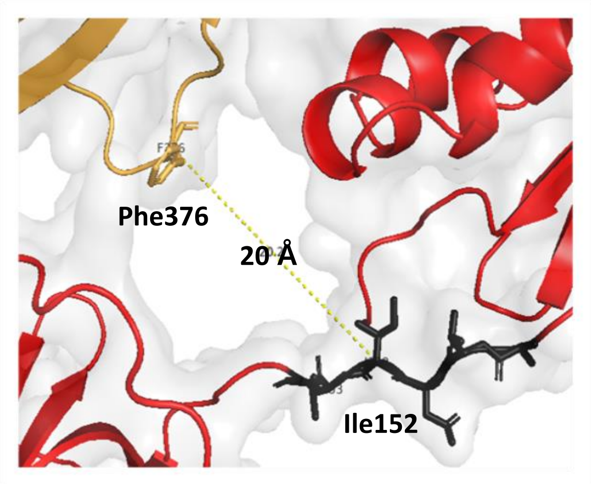


**B**

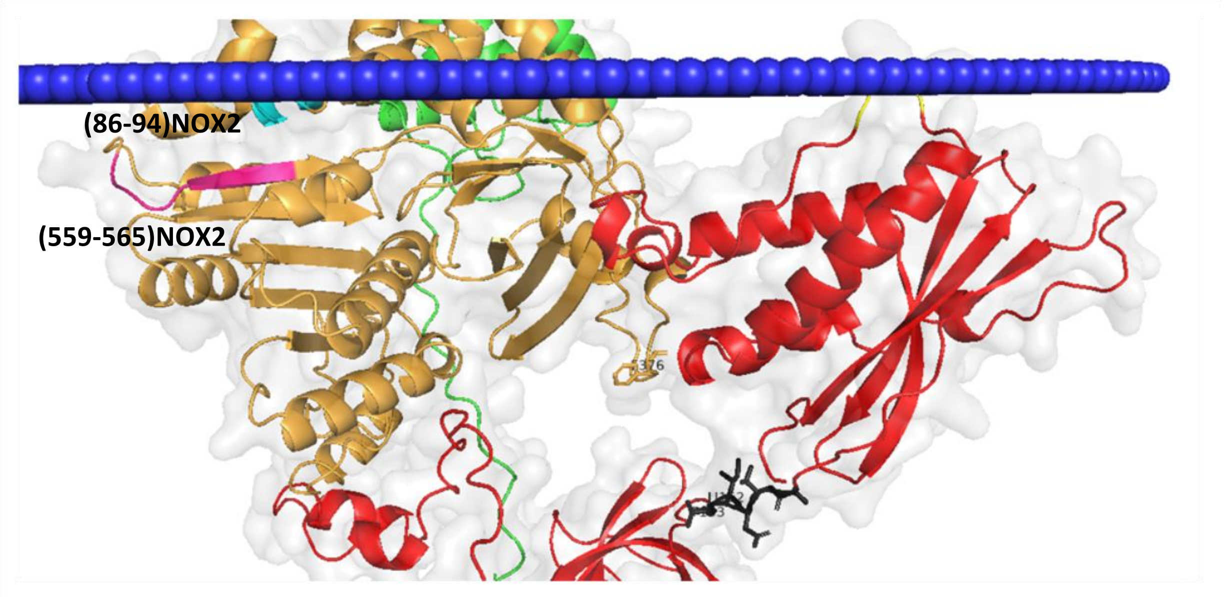


**Figure S4. A.** Distance between residue **Ile152 of p47$^{phox}$ and residue Phe376 of NOX2**. **B. Region 86-94 of NOX2** (cyan) located in loop B and region **559-565 of NOX2** C-terminus (colored in magenta). These two regions of NOX2 (colored in light brown) are structurally close and are positioned far from p47$^{phox}$ (red) in this model. In green is colored p22$^{phox}$.

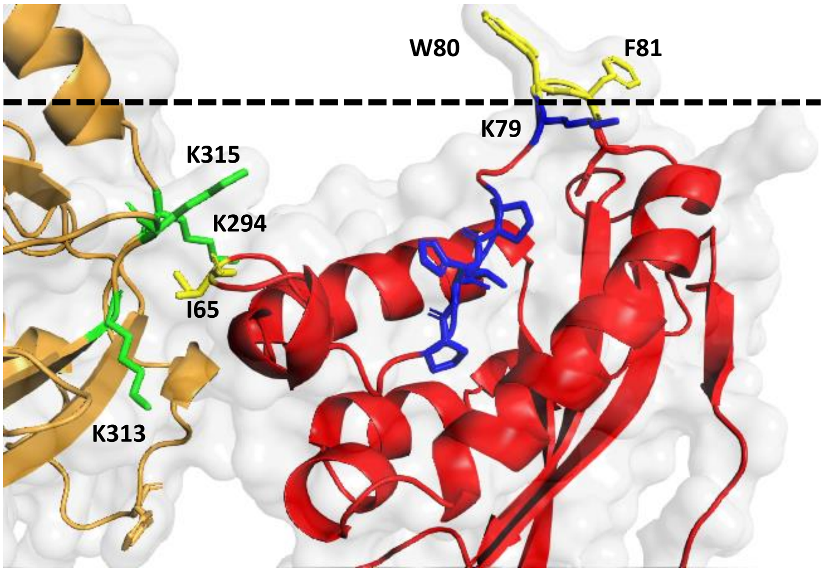


**Figure S5. Interacting regions of p47$^{phox}$ with membrane.** NOX2 and p47$^{phox}$ are colored in light brown and red, respectively. The Ile65 previously proposed as the site of p47phox insertion into the membrane is found in our model in a pocket formed by three lysines (K294, K313, K315). Insertion into the membrane occurs instead at aromatic residues (W80 and Phe81). The critical residues involved in canonical recognition of phosphoinositides (PtdIns3P) “RPPR” (R43,P73,P76,R90) are colored in blue. The predicted membrane calculated with PPM has been removed for a better view and is symbolized here by the dotted black line representing the level of the polar head groups of the phospholipids in the inner leaflet of the membrane.

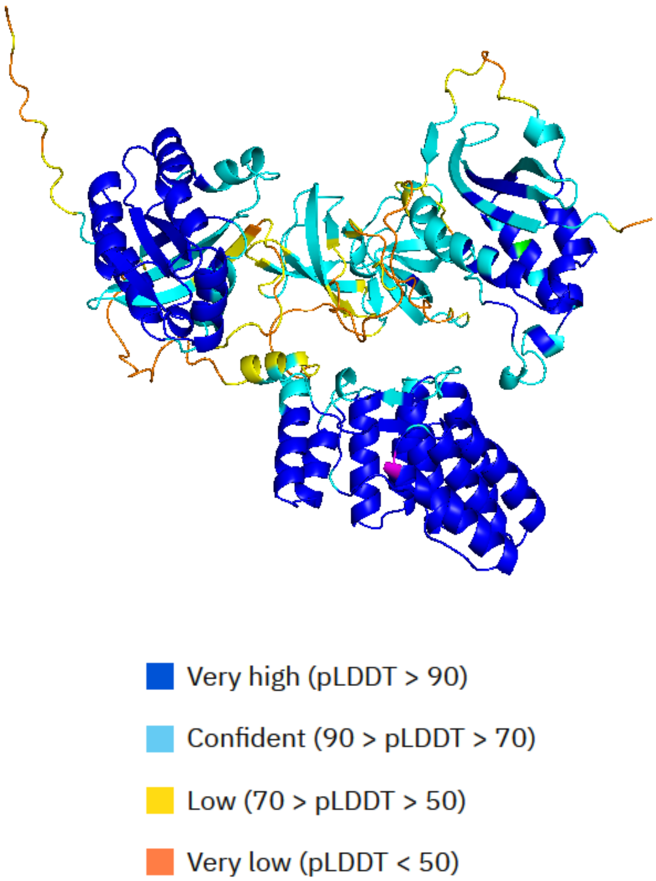


**Figure S6. AlphaFold model of the Trimera**Staining of the model in pLLDT suggests a high degree of confidence for the $p67^{phox}$ protein and Rac1 in large part and a lower degree for the $p47^{phox}$ protein.

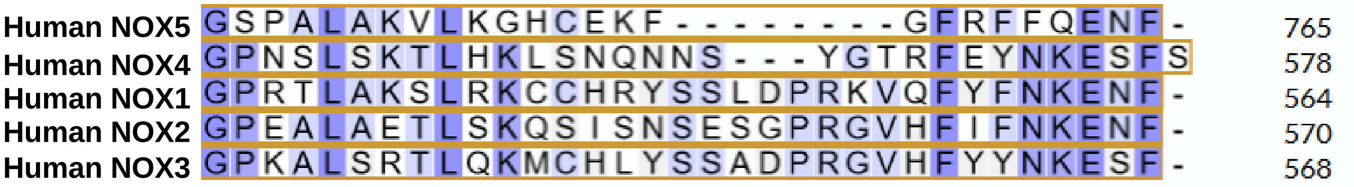


**Figure S7. BLAST of human NOX1, NOX2, NOX3, NOX4 and NOX5 showing sequence similarity at their C-terminus.** The presence of a serine as terminal residue is present only in constitutively active NOX4. All other activatable NOX proteins (1, 2, 3 and 5) end with Phenylalanine.
